# Hyperactivation of the AXL-ICD/SIRT2 axis by Amyloid-β impairs astrocytic autophagic flux and exacerbates neuroinflammation

**DOI:** 10.64898/2026.08.23.746508

**Authors:** Tai Young Kim, Mridula Bhalla, Uiyeol Park, Seung Jae Hyeon, In-Young Hwang, Youngsuk Seo, Wongu Youn, Jin-A Lee, Junghee Lee, Boyoung Lee, Hoon Ryu, C. Justin Lee

## Abstract

Autophagy dysfunction and neuroinflammation are central to Alzheimer’s disease (AD), yet how extracellular amyloid-β (Aβ) couples to impaired autophagic flux and heightened neuroinflammation remains unknown. Here, we identify the TAM receptor AXL as a molecular transducer that couples Aβ sensing to the regulation of autophagy and neuroinflammation in astrocytes. Aβ induces γ-secretase-dependent cleavage of AXL, generating a nuclear intracellular domain (AXL-ICD) that forms phase-separated condensates and activates autophagy gene transcription through SIRT2-mediated recruitment of the RUVBL1/2-INO80 chromatin-remodeling complex. This axis is activated in astrocytes of postmortem AD brains. Concurrently, AXL-ICD binds to the SIRT2 catalytic domain and suppresses its deacetylase activity, increasing α-tubulin acetylation and altering microtubule dynamics. While moderate AXL-ICD levels promote autophagic flux, excessive elevation paradoxically triggers microtubule hyperstabilization, thereby impairing autophagosome-lysosome fusion and causing pathological accumulation of autophagosomes and H_2_O_2_. The inhibitory peptide AxSBiP disrupts the AXL-ICD/SIRT2 interaction, restores autophagic flux, reduces plaque burden, and normalizes Aβ-induced H_2_O_2_ production and astrogliosis in APP/PS1 mice. We propose the AXL-ICD/SIRT2 axis as an effective therapeutic target to reduce Aβ burden and neuroinflammation in AD.

## Introduction

The clinical limitations of current monoclonal antibody therapies targeting Aβ in AD underscore the need for alternative therapeutic approaches. A recent systematic review concluded that Aβ removal by monoclonal antibodies (e.g., aducanumab, bapineuzumab, crenezumab, donanemab, gantenerumab, lecanemab, ponezumab, remternetug, and solanezumab) does not consistently translate into clinically meaningful cognitive improvement^1^. More serious concerns include the significant risk of amyloid-related imaging abnormalities (ARIA), which reflect treatment-induced neurovascular perturbations and neuroinflammation, highlighting the need to explore alternative disease-modifying mechanisms beyond Aβ clearance.

Autophagy is a fundamental cellular process that mediates the clearance of protein aggregates and other intracellular components in response to various stressors, such as nutrient deprivation, thereby maintaining proteostasis^2,3^. While canonical pathways governing autophagy induction, primarily mediated by posttranslational modifications, are well established^4,5^, how extracellular protein aggregates are sensed and coupled to the transcriptional activation of autophagy genes, and how this linkage relates to neuroinflammation, remain elusive. Notably, dysregulation of autophagy has been widely reported in AD, where accumulation of autophagic vacuole is interpreted as a consequence of impaired autophagic flux rather than increased autophagosome formation *per se*^6,7^. More importantly, we have recently reported that impaired autophagic flux along with astrogliosis in astrocytes is the hallmark of AD^8^. These observations reflect an uncoupling between autophagy induction and degradation in AD. Accordingly, how disease-associated extracellular signals perturb the regulation of autophagy and neuroinflammation and contribute to this uncoupling remains a critical question for preserving proteostasis and preventing neuroinflammation.

The TAM receptor family, comprising TYRO3, AXL, and MERTK, represents a candidate link between extracellular cues and the regulation of autophagy. These receptors facilitate phagocytic clearance of extracellular substrates, including apoptotic cells and myelin debris^9^. In the brain, AXL and MERTK have been implicated in the detection and clearance of Aβ plaques, with astrocytes particularly contributing to intracellular Aβ degradation^10–13^. Although AXL has been implicated in autophagy regulation through kinase-dependent cytoplasmic signaling^14,15^, whether it links extracellular Aβ sensing to kinase-independent autophagy regulation remains unclear.

SIRT2, a member of the sirtuin family that functions as a class III histone deacetylase (HDAC), plays well-established roles in cell cycle regulation and cytoskeletal maintenance, particularly through deacetylation of α-tubulin^16,17^. In neuronal models of AD, SIRT2-mediated α-tubulin deacetylation has been shown to impair microtubule dynamics and autophagic flux^18–20^. In our recent studies on AD, we identified cytosolic SIRT2 as the most highly expressed HDAC in astrocytes and demonstrated its critical role in the astrocytic cytoplasmic putrescine degradation pathway, leading to MAOB-dependent production of GABA, H_2_O_2_, and ammonia^21,22^, which are tightly linked to neuroinflammation, memory impairment, and neurodegeneration in AD^23^. Despite these advances, whether SIRT2 contributes to nuclear transcriptional induction of autophagy and whether SIRT2 inhibition-driven α-tubulin hyperacetylation impairs microtubule dynamics and autophagosome trafficking in astrocytes under AD conditions remain entirely unexplored.

Here we show that Aβ induces γ-secretase-dependent cleavage of AXL to generate AXL-ICD, which co-translocates with SIRT2 into the nucleus to recruit chromatin-remodeling machinery and activate autophagy gene transcription. Concurrently, AXL-ICD suppresses SIRT2 deacetylase activity on α-tubulin in a dose-dependent manner: while moderate activation promotes autophagic flux, excessive AXL-ICD accumulation causes microtubule hyperstabilization, failure of autophagosome-lysosome fusion, and over-production of H_2_O_2_. Disrupting this axis with the competitive inhibitory peptide AxSBiP restores autophagic flux, normalizes H_2_O_2_ production and astrogliosis, and reduces Aβ plaque burden in APP/PS1 mice, identifying the AXL-ICD/SIRT2 interface as a therapeutic target for AD.

## Results

### AXL functions as an Aβ receptor that induces autophagy gene expression in astrocytes

We have previously shown that Aβ induces autophagy-related genes in astrocytes^8^. Consistently, Aβ treatment markedly increased LC3B, p62/SQSTM1 and ATG5 in human astrocytes (Supplementary Fig. 1a, b). Because TAM receptors have recently been identified as Aβ receptors^11^ and are highly expressed in astrocytes (Supplementary Fig. 1c), we tested whether they link Aβ signaling to autophagy gene induction. Silencing AXL, but not other TAM receptors, markedly reduced LC3B protein levels (Supplementary Fig. 1d-f), accompanied by decreased *LC3B* mRNA levels (Supplementary Fig. 1g), indicating transcriptional regulation. Consistently, AXL depletion reduced autophagic flux, as evidenced by diminished LC3-II and p62 accumulation after bafilomycin A1 treatment (Supplementary Fig. 1h). Notably, AXL silencing abolished Aβ-induced LC3B upregulation (Fig. 1a, b). Together, these findings establish AXL as a critical receptor that links extracellular Aβ sensing to transcriptional activation of the autophagy genes in astrocytes.

**Fig 1.**
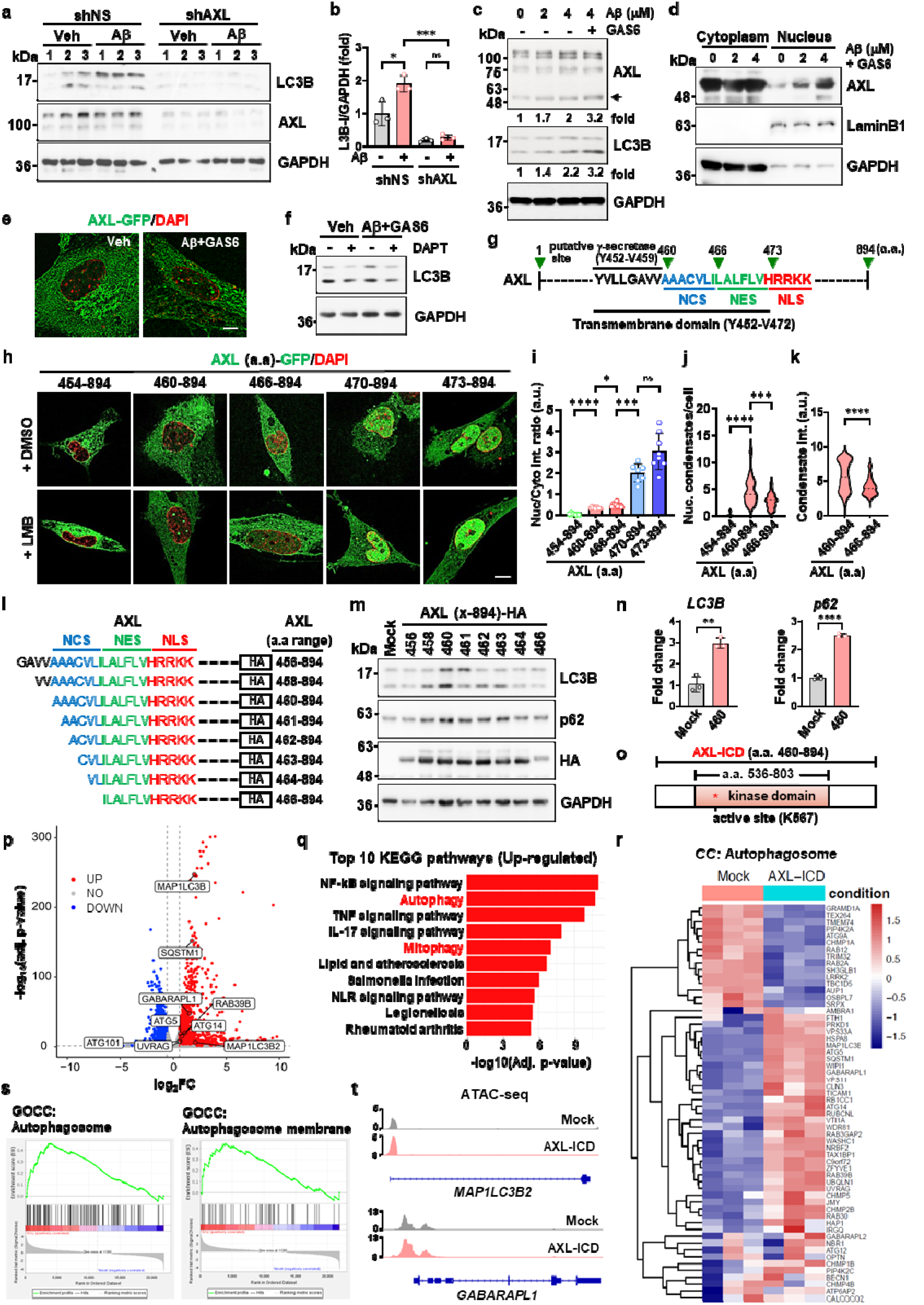
Aβ induces AXL cleavage and nuclear AXL-ICD to regulate autophagy gene transcription in astrocytes. **a, b** Immunoblots and quantification of LC3B-I levels in Aβ oligomers treated astrocytes (4 µM, 24 h) with or without AXL knockdown. **c** Dose-dependent effects of Aβ oligomers on AXL cleavage and LC3B levels in the presence or absence of GAS6 (250 ng/ml). Arrow indicates AXL-ICD. **d** Immunoblots of AXL in cytoplasmic and nuclear fractions following Aβ and GAS6 (250 ng/ml) treatment. **e** Representative images of C-terminal GFP-tagged AXL localization in astrocytes treated with 4 µM Aβ and 250 ng/ml GAS6. **f** Effect of γ-secretase inhibitor DAPT (10 µM) on Aβ-induced LC3B levels. **g** Schematic of AXL transmembrane region highlighting the γ-secretase cleavage site and nuclear signaling motifs (NLS, NCS, and NES). **h, i** Localization of AXL deletion constructs and nuclear-to-cytosolic intensity ratios in astrocytes treated with Leptomycin B (LMB, 10 nM). **j, k** Quantification of nuclear condensate number and intensity for indicated AXL constructs in LMB-treated astrocytes. **l** Schematic of HA-tagged AXL fragments. Colored residues indicate NCS (blue), NES ^35^, and NLS (red). Amino acid ranges are indicated on the right. **m** Immunoblots of LC3B and p62 in U87MG cells expressing HA-tagged AXL deletion constructs. **n** qRT-PCR analysis of *LC3B* and p62 mRNA normalized to GAPDH in AXL-ICD-overexpressing U87MG cells. (*n* = 3). **o** Schematic of AXL-ICD. The kinase domain (aa 536–803) and the catalytic lysine residue (K567, asterisk) are indicated. **p, q** Volcano plot and KEGG pathway enrichment analysis of upregulated genes in AXL-ICD-overexpressing U87MG cells. **r** Heatmap of autophagosome-related genes (GO:0005776) using FPKM-normalized expression values. **s** Gene ontology enrichment analysis (GSEA) plots showing significant enrichment of “autophagosome” and “autophagosome membrane” gene sets. **t** ATAC-seq tracks showing chromatin accessibility at *MAP1LC3B2* and *GABARAPL1* loci. Data are mean ± s.e.m. Statistical significance was determined by two-way ANOVA with Sidak’s multiple-comparison test (**b**), one-way ANOVA with Tukey’s multiple-comparison test (**i, j**), or unpaired two-tailed Student’s t-test (**k, n**). ns, not significant; *P < 0.05, **P < 0.01, ***P < 0.001, ****P < 0.0001.

### Aβ-induced γ-secretase cleavage of AXL generates a nuclear AXL intracellular fragment

We next investigated how Aβ induces autophagy-related gene expression through AXL. AXL undergoes sequential α- and γ-secretase cleavage to generate an intracellular domain (AXL-ICD) capable of nuclear translocation^24,25^, although its function remains unknown. We therefore asked whether Aβ promotes AXL cleavage and nuclear localization of AXL-ICD to regulate autophagy-related gene expression. In human astrocytes, Aβ induced accumulation of ∼50-kDa AXL fragments corresponding to AXL-ICD without noticeable alteration in total AXL levels, and this processing was further enhanced by GAS6, a high-affinity AXL ligand ^26^ (Fig. 1c). A similar but more pronounced increase in AXL-ICD accumulation was observed in U87MG astrocytoma cells (Supplementary Fig. 1i). Cellular fractionation revealed a dose-dependent increase in nuclear AXL-ICD with increasing Aβ in the presence of GAS6 (Fig. 1d). Co-treatment induced nuclear GFP signals from C-terminally tagged AXL, indicating nuclear localization of the C-terminal AXL-ICD that formed condensate-like structures (Figure 1E), which was further confirmed by immunostaining with a C-terminal-specific AXL antibody (Supplementary Fig. 1j). Finally, the γ-secretase inhibitor DAPT blocked the Aβ-induced increase in LC3B (Fig. 1f), suggesting that γ-secretase-dependent AXL cleavage contributes to autophagy gene induction. Together, these results indicate that Aβ promotes γ-secretase-dependent cleavage of the AXL receptor to generate nuclear AXL-ICD that drives autophagy-related gene expression.

A putative γ-secretase cleavage site of AXL has been mapped to amino acids (aa) 452 to 459 within the transmembrane region (aa 452–472)^25^ (Fig. 1g), although the exact cleavage site may vary within this region owing to the broad sequence specificity of γ-secretase^27^. To define the exact AXL region responsible for nuclear translocation and autophagy gene induction, we generated a series of AXL deletion constructs with a C-terminal GFP tag (Supplementary Fig. 2a). Constructs retaining most of the transmembrane domain (aa 454– 894) failed to show nuclear GFP signals even after treatment with leptomycin B (LMB; a potent nuclear export inhibitor) (Fig. 1h, i), indicating membrane retention. By contrast, the aa 460–894 construct exhibited enhanced nuclear GFP signals that formed nuclear condensates upon LMB treatment (Fig. 1h, i), resembling those observed in Aβ-treated cells (Fig. 1e). Further truncation (aa 466–894) increased nuclear localization but reduced condensate formation (Fig. 1h-k), indicating that residues 460–465 are required for nuclear condensate formation and define a previously unrecognized nuclear condensate sequence (NCS) (Fig. 1g). Constructs lacking residues 466–469 (aa 470–894 and aa 473–894), showed robust nuclear localization independent of LMB treatment (Fig. 1h, i), consistent with a predicted nuclear localization signal (NLS) at residues 473–477 (HRRKK)^25^. Notably, residues 466–473 suppressed this nuclear localization (Fig. 1h, i), suggesting nuclear export activity within this region. Supporting this, residues 465–472 (LILALFLV) harbor a hydrophobic sequence resembling a classical nuclear export signal (NES) (Supplementary Fig. 2b). Similar patterns were observed in U87MG cells (Supplementary Fig. 2c,d). Given AXL aa 460–894 fragment formed nuclear condensates, we next examined their biophysical properties. Live-cell imaging showed that AXL-ICD-GFP formed dynamic, spherical nuclear condensates that underwent fusion and fission (Supplementary Fig.2e), and were rapidly dissolved by 1,6-hexanediol (1,6-HD) (Supplementary Fig. 2f, g), indicating liquid-liquid phase separation (LLPS)-like properties^28^ They showed minimal overlap with the nucleolar marker NPM1 (Supplementary Fig. 2h), indicating that AXL aa 460–894 fragment forms non-nucleolar nuclear condensates. Together, these results indicate that AXL aa 460–894 fragment contains distinct NCS, NES, and NLS sequences that act in concert to regulate nuclear entry and condensate formation, and forms dynamic, non-nucleolar nuclear condensates through LLPS

### AXL-ICD drives autophagy gene transcription and autophagosome formation

Based on the above characterization of the AXL aa 460–894 fragment, we generated a series of deletion variants centered on residue 460 (Fig. 1l) and assessed their ability to induce autophagy genes including *LC3B* and *p62* in U87MG cells. These cells exhibit more robust and reproducible responses than human astrocytes, enabling reliable assessment of AXL-ICD-dependent autophagy activation. Notably, the aa 460–894 fragment robustly upregulated both LC3B and p62, whereas deletion aa 460–463 (AAAC; hereafter “AAAC motif”) within the NCS completely abolished this activity (Fig. 1m). Furthermore, even minimal N-terminal extensions (aa 456–894 or aa 458–894) markedly attenuated LC3B and p62 induction (Fig. 1m), highlighting a sharp positional requirement at residue 460. Importantly, qRT-PCR analysis confirmed that the aa 460–894 fragment transcriptionally upregulated *LC3B* and *p62* mRNA levels (Fig. 1n). Based on these findings, we define the aa 460–894 fragment as AXL-ICD (Fig. 1o), which is generated by Aβ-induced γ-secretase cleavage (Fig. 1c). Notably, this cleavage site shares sequence similarity with the γ-secretase cleavage site of amyloid precursor protein at position 40 (Supplementary Fig. 3a). Supporting this definition, AXL-ICD overexpression recapitulated the Aβ-induced autophagy-related expression profile, increasing LC3B, p62 and ATG5 (Supplementary Fig. 3b, c, and 1b). In addition, it selectively upregulated GABARAPL1, but not GABARAP, two ATG8-family members, which is associated with cargo recruitment and late-stage autophagosome maturation^29^ (Supplementary Fig. 3d, e).

To examine the broader transcriptional impact of AXL-ICD, we performed RNA sequencing in U87MG cells overexpressing AXL-ICD. Differential expression analysis revealed robust upregulation of autophagy genes, including *MAP1LC3B, MAP1LC3B2, SQSTM1* and *GABARAPL1* (Fig. 1p). Pathway enrichment analysis using the KEGG database showed that autophagy and mitophagy were among the top-enriched pathways, along with inflammation-related pathways including NF-κB, TNF, and IL-17 signaling (Fig. 1q, Supplementary Fig. 4a). Heatmap clustering further revealed coordinated induction of autophagy-related gene signatures in AXL-ICD-overexpressing cells (Fig. 1r). Consistently, Gene Set Enrichment Analysis demonstrated significant enrichment of gene sets associated with autophagosome organization and membrane components (Supplementary Fig. 1s). Notably, this transcriptional program was distinct from canonical TFEB-driven lysosomal biogenesis, as AXL-ICD neither interacted with TFEB nor induced lysosomal gene expression (Supplementary Fig. 4b, c). To assess changes in chromatin accessibility associated with autophagy gene activation, we performed Assay for Transposase-Accessible Chromatin using sequencing (ATAC-seq). AXL-ICD overexpression increased promoter accessibility at key autophagy-related genes, including the ATG8 family members *MAP1LC3B2* and *GABARAPL1* (Fig. 1t), indicating enhanced chromatin accessibility at autophagy gene promoters.

To explore the structural basis of AXL-ICD, we performed structural modeling, which predicted a dimeric configuration (Supplementary Fig. 5a), supported by co-immunoprecipitation of HA- and FLAG-tagged constructs (Supplementary Fig. 5b). Importantly, both this interaction and the induction of LC3B and p62 were independent of AXL kinase activity, as the kinase-dead K567R mutant showed effects comparable to wild-type (Supplementary Fig. 5b-d). Finally, to investigate whether the upregulation of autophagy-related proteins by AXL-ICD is directly linked to autophagosome formation, we employed both Cyto-ID staining and autophagy-specific probes. First, flow cytometric analysis using the Cyto-ID dye^30^ revealed that AXL-ICD expression led to an approximately 1.5-fold increase in autophagic compartments (Figure S5E). Second, using autophagy-specific probes with LIR motifs targeting distinct mATG8 subfamilies including FYCO1 for LC3A/LC3B and ATG4B(T) for GABARAP/GABARAPL1^31^, AXL-ICD expression significantly increased autophagic puncta, indicating enhanced autophagosome formation (Supplementary Fig. 5f, g). Together, these findings identify AXL-ICD as an effector of Aβ signaling that drives autophagy gene transcription via enhanced chromatin accessibility at autophagy gene promoters, ultimately promoting autophagosome formation.

### AXL-ICD hijacks SIRT2 to couple autophagy gene transcription with cytoplasmic microtubule remodeling

Because AXL-ICD lacks recognizable transcriptional or epigenetic regulatory motifs, we hypothesized that it functions as a platform to recruit nuclear cofactors. Given that SIRT2 translocates to the nucleus under pathological conditions^21,32–34^ and is highly expressed in astrocytes and implicated in AD-related pathways^21,23^, we tested whether it serves as a nuclear partner of AXL-ICD in astrocytes. Aβ treatment induced nuclear accumulation of SIRT2 in primary mouse astrocytes and U87MG cells (Supplementary Fig. 6a and Fig 2a, b). This effect was markedly attenuated by AXL silencing, which also reduced basal nuclear SIRT2 levels, indicating that AXL contributes to both basal and Aβ-induced SIRT2 nuclear localization (Fig. 2a, b). Importantly, AXL-ICD overexpression alone was sufficient to drive SIRT2 nuclear accumulation even in the absence of Aβ (Fig. 2c, d). Moreover, this effect was dose-dependent, correlating with nuclear AXL-ICD levels in a doxycycline-inducible system (Supplementary Fig. 6b), suggesting that AXL-ICD acts as a primary driver of SIRT2 nuclear translocation. Immunoprecipitation analyses revealed that AXL-ICD preferentially interacts with SIRT2 among sirtuin family proteins (Fig. 2e and Supplementary Fig. 5c), and proximity ligation assays^35^ confirmed robust AXL-ICD/SIRT2 association in both nuclear and cytoplasmic compartments (Fig. 2f-h). This interaction occurred independently of enzymatic activity, as kinase-inactive AXL and deacetylase-deficient SIRT2 mutants retained binding (Supplementary Fig. 6d, e). To delineate the protein-protein interaction interface between these two proteins, we performed deletion mapping analysis, revealing that the AAAC motif in AXL-ICD (aa 460–463) (Supplementary Fig. 6f and 2i) and the catalytic domain of SIRT2 (Supplementary Fig. 6g and 2j) are critical for their association. Strikingly, this AAAC motif coincides with the NCS required for condensate formation (Fig. 1h and 1j) and autophagy gene induction (Fig. 1l, m). Structural modeling further predicted that the AAAC motif of AXL-ICD directly engages the catalytic domain of SIRT2 (Fig. 2k), establishing a structural basis for this interaction. These findings led us to hypothesize that SIRT2 is an essential cofactor for AXL-ICD-dependent autophagy regulation. Consistent with this, SIRT2 depletion markedly suppressed the AXL-ICD-mediated induction of LC3B and p62 (Fig. 2l, m). This defective autophagy response was associated with a significant reduction in AXL-ICD nuclear condensates (Supplementary Fig. 6h, i). Together with the observation that AGK2-mediated inhibition of SIRT2 catalytic activity did not alter AXL-ICD-mediated LC3B or p62 induction (Supplementary Fig. 6j, k), these results indicate that SIRT2 supports AXL-ICD-driven autophagic signaling through a non-enzymatic, structural mechanism rather than its deacetylase activity.

**Fig. 2.**
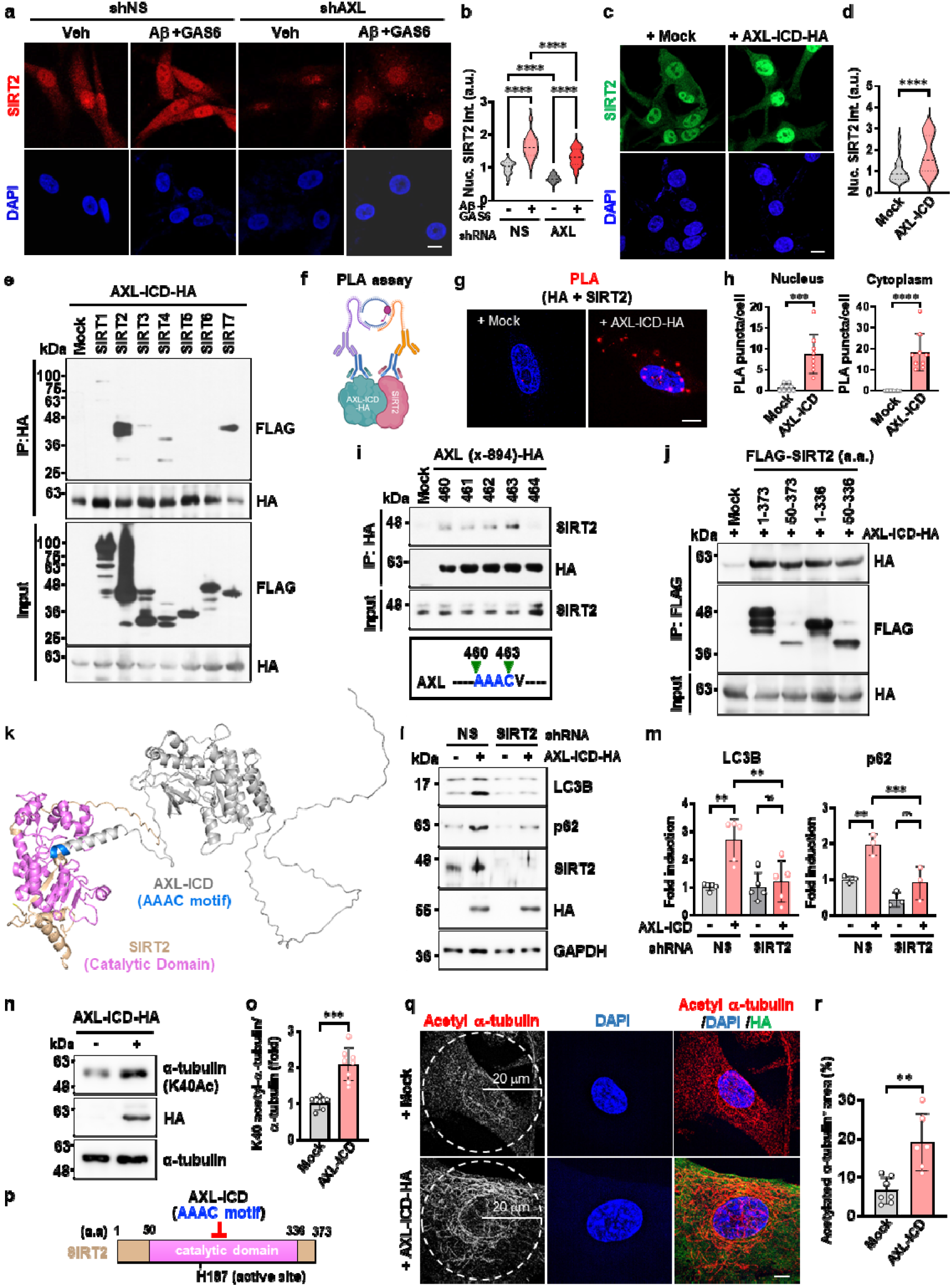
AXL-ICD interacts with SIRT2 to promote its nuclear localization and suppress deacetylase activity. **a, b** Representative images and quantification of nuclear SIRT2 intensity in U87MG cells treated with 4 µM Aβ and 250ng/ml GAS6 with or without AXL knockdown. (*n* = 35-52 cells/group). Scale bars, 10 μm. **c, d** Representative images and quantification of nuclear SIRT2 intensity in AXL-ICD-overexpressing U87MG cells. (*n* = 44-48 cells/group). Scale bars, 10 μm. **e** Co-immunoprecipitation of FLAG-tagged sirtuin proteins with HA-tagged AXL-ICD in HEK293T cells. **f-h** Schematic of Proximity Ligation Assay^35^ (**f**), representative PLA signals (**g**), and quantification of nuclear and cytoplasmic PLA puncta per cell (*n* = 9) (**h**). Scale bar, 5 μm. **i** Co-immunoprecipitation of endogenous SIRT2 with HA-tagged AXL-ICD deletion constructs in U87MG cells. (Bottom) Minimal SIRT2-binding AAAC motif (aa 460–463). **j** Co-immunoprecipitation of SIRT2 deletion mutants with HA-tagged AXL-ICD constructs in U87MG cells. **k** Structural model of the interaction between AXL-ICD and SIRT2 generated using the Schrödinger modeling suite and visualized in PyMOL. **l, m** Immunoblots and quantification of LC3B (*n* = 5) and p62 (*n =* 3) in AXL-ICD-overexpressing U87MG cells with or without SIRT2 knockdown. **n, o** Western blots and quantification of acetylated α-tubulin (K40) in AXL-ICD-overexpressing U87MG cells. (*n* = 6-8 /group). **p** Proposed model illustrating the interaction between the AXL-ICD AAAC motif and the SIRT2 catalytic domain, leading to enzymatic inhibition. **q, r** Immunofluorescence) and area-based quantification of acetylated α-tubulin in the perinuclear region. (*n* = 6-7 cells /group). Scale bar, 5 μm. Data are mean ± s.e.m. Statistical significance was determined by two-way ANOVA with Tukey’s post hoc test (**b, m**) or unpaired two-tailed Student’s t-test (**d, h, q,** and **r**). ns, not significant; **P < 0.01, ***P < 0.001, ****P < 0.0001.

Although AXL-ICD-driven autophagic signaling occurs independently of SIRT2 catalytic activity, AXL-ICD was found to bind the catalytic domain of SIRT2, raising the question of whether this interaction might instead inhibit SIRT2 deacetylase activity. Intriguingly, AXL-ICD expression increased acetylation of α-tubulin at lysine 40 (K40), a well-established deacetylation site on α-tubulin regulated by SIRT2^36^ (Fig. 2n, o), indicating suppression of SIRT2 deacetylase activity (Fig. 2p). Consistently, the acetylated α-tubulin⁺ area was markedly increased (Fig 2q, r), indicative of enhanced microtubule stabilization, which has been linked to efficient autophagosome–lysosome fusion and autophagic flux^37^. To further support SIRT2 inhibition by AXL-ICD, transcriptomic analysis revealed downregulation of cell cycle-related gene programs, resembling expression profiles observed following SIRT2 inhibition or depletion^16,38,39^ (Supplementary Fig. 6l, m). Together, these findings indicate that AXL-ICD utilizes SIRT2 as a nuclear scaffold to promote condensate assembly and autophagy gene activation, while simultaneously suppressing its deacetylase activity to stabilize cytoplasmic microtubules. This dual mechanism effectively links nuclear transcriptional activation to cytoplasmic autophagy regulation.

### AXL-ICD/SIRT2 recruits the RUVBL1/2/INO80 complex to activate autophagy genes

Given that AXL-ICD binds SIRT2 thereby suppressing its catalytic activity, we postulated that, following complex formation with SIRT2, AXL-ICD recruits additional cofactors to drive autophagy gene activation. To identify such factors, we performed immunoprecipitation followed by mass spectrometry in human astrocytes and U87MG cells overexpressing AXL-ICD-HA (Fig. 3a). The proteomics analysis identified several AXL-ICD-interacting proteins in both cell lines (Fig. 3b). Top-ranked interactors included transcriptional regulators such as RUVBL1, RUVBL2, and WDR5, as well as autophagy-related factors including ERLIN1/2 and ATAD3A (Fig. 3a and Supplementary Fig. 7a). These interactions were validated by immunoprecipitation and western blotting analyses (Supplementary Fig. 7b). Gene ontology analysis of AXL-ICD-associated proteins revealed enrichment for organelle organization, protein folding, and protein metabolic processes, but not canonical autophagy components (Supplementary Fig. 7c), further supporting a transcriptional mode of AXL-ICD-mediated autophagy regulation. Among these interactors, we focused on RUVBL1 and RUVBL2 (Pontin and Reptin), ATPases that function as core components of chromatin-remodeling and transcriptional regulatory complexes^40,41^. Notably, RUVBL1/2 have been implicated in autophagy regulation^42^ and are enriched in the SIRT2 interactome^43^. PLA revealed associations of AXL-ICD with RUVBL1 and RUVBL2 in both nuclear and cytoplasmic compartments (Fig. 3c, d). Consistent with prior work ^43^, RUVBL1/2 also interacted with SIRT2 (Fig. 3e), and SIRT2 overexpression enhanced, whereas its depletion reduced, AXL-ICD/RUVBL1 association (Fig. 3f-h), indicating that SIRT2 mediates the recruitment of the RUVBL1/2 complex to AXL-ICD. Functional analyses demonstrated that shRNA-mediated silencing of both RUVBL1 and RUVBL2 abolished AXL-ICD-induced increases in LC3B and p62 protein levels (Fig. 3i, j). Similarly, pharmacological inhibition of RUVBL1/2 with CB-6644 dose-dependently reduced AXL-ICD-induced LC3B and p62 expression (Supplementary Fig. 8a, b). These results indicate that the RUVBL1/2 complex is required for AXL-ICD/SIRT2-dependent activation of autophagy gene transcription.

**Fig. 3.**
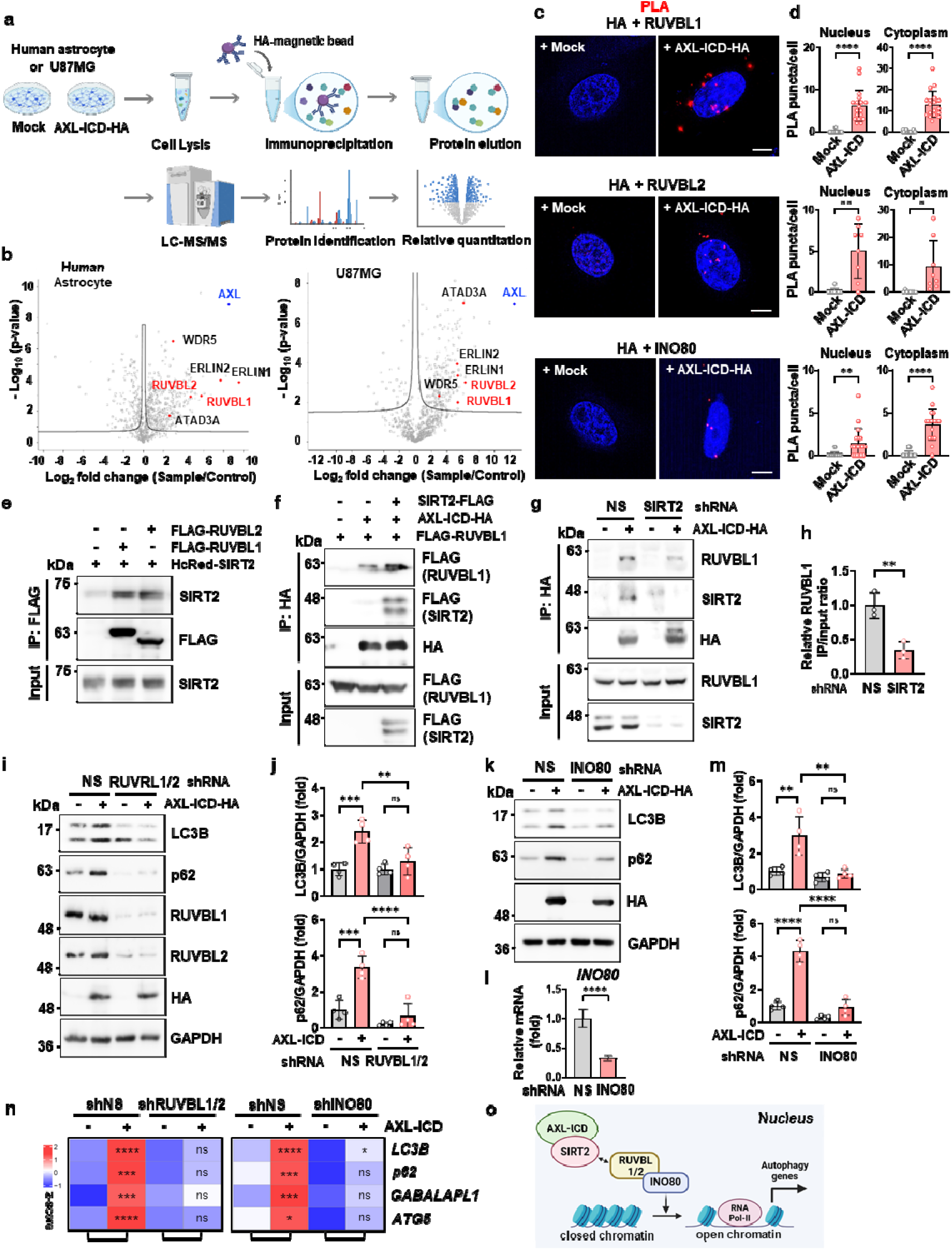
Identification of AXL-ICD-interacting proteins involved in autophagy regulation. **a** Schematic of the experimental workflow. Proteomic approach using anti-HA immunoprecipitation and LC-MS/MS to identify AXL-ICD-interacting proteins in human astrocytes and U87MG cells. **b** Volcano plots of AXL-ICD interactors. Proteins enriched in AXL-ICD-HA immunoprecipitates (FDR < 0.05) are shown. AXL (blue) and selected interactors (RUVBL1/2, WDR5, ERLIN1/2, and ATAD3A) are highlighted. **c, d** PLA analysis of AXL-ICD and chromatin-remodeling factors. Representative PLA signals and quantification of puncta per cell showing proximity between AXL-ICD and endogenous RUVBL1 (*n* = 18 cells), RUVBL2 (*n* = 8 cells), or INO80 (*n* = 19 cells). Scale bar, 5 μm. **e** Co-immunoprecipitation of FLAG-tagged RUVBL1 or RUVBL2 with HcRed-tagged SIRT2 in HEK293T cells. **f** Co-immunoprecipitation of FLAG-tagged RUVBL1 with HA-tagged AXL-ICD in HEK293T cells in the presence or absence of FLAG-tagged SIRT2. **g, h** Co-immunoprecipitation of endogenous RUVBL1 with HA-tagged AXL-ICD in U87MG cells expressing shNS or shSIRT2, with quantification of co-immunoprecipitated RUVBL1 levels (*n* = 3). **i, j** Immunoblots and quantification of LC3B and p62 in AXL-ICD-overexpressing U87MG cells with or without RUVBL1/2 knockdown. (*n* = 4). **k-m** Immunoblots of LC3B and p62 (**k**), qRT-PCR validation of INO80 knockdown (**l**), and quantification of LC3B and p62 levels (*n* = 4) (**m**) in AXL-ICD-expressing U87MG cells with or without INO80 knockdown. **n** qRT-PCR analysis showing relative mRNA levels of *LC3B, p62, GABARAPL1*, and *ATG5* in AXL-ICD-expressing U87MG cells with or without knockdown of RUVBL1/2 (left) or INO80 (right). Data are presented as a z-score heatmap (*n* = 3). **o** Proposed model of the AXL-ICD/SIRT2/RUVBL/INO80 complex. Schematic illustrating the recruitment of the RUVBL1/2-INO80 chromatin-remodeling complex by AXL-ICD/SIRT2 to promote the transcription of autophagy-related genes. Data are presented as mean ± s.e.m. Statistical significance was determined by unpaired two-tailed Student’s t-test (**d, h**) or two-way ANOVA with Tukey’s post hoc test (**j, m**). ns, not significant; *P < 0.05, **P < 0.01, ***P < 0.001, ****P < 0.0001.

Previous studies have shown that under glucose starvation, CARM1-mediated methylation of RUVBL1 promotes recruitment of the TIP60 chromatin-remodeling complex to autophagy gene loci^42^. To determine whether the AXL-ICD/SIRT2/RUVBL1/2 pathway operates through a similar mechanism, we examined the roles of CARM1 and TIP60. Genetic and pharmacological inhibition of CARM1 and TIP60 did not affect AXL-ICD-induced LC3B and p62 expression (Supplementary Fig. 8c-g), indicating that this pathway operates independently of CARM1/TIP60 signaling. Because RUVBL1/2 participate in multiple chromatin-remodeling complexes, including TIP60 and INO80^44^, we next examined the involvement of INO80. Targeted immunoprecipitation further revealed an interaction between AXL-ICD and INO80, despite its absence from the proteomic dataset (Supplementary Fig. 8h), and PLA assays detected AXL-ICD/INO80 association in both nuclear and cytoplasmic compartments (Fig. 3c, d, bottom panels). Silencing INO80 abolished AXL-ICD-induced upregulation of LC3B and p62 proteins (Fig. 3k-m) and similarly reduced autophagosome formation as measured by Cyto-ID staining (Supplementary Fig. 8i, j). Critically, qRT-PCR analysis demonstrated that silencing of RUVBL1/2 or INO80 suppressed transcriptional induction of *LC3B*, *p62*, *GABARAPL1*, and *ATG5* (Fig. 3n), directly confirming that AXL-ICD-mediated autophagy regulation occurs at the transcriptional level through the RUVBL1/2/INO80 complex. Collectively, these findings establish a receptor-driven epigenetic mechanism whereby AXL-ICD activates autophagy gene transcription through SIRT2-dependent recruitment of chromatin-remodeling machinery, distinct from the starvation-induced CARM1/TIP60 pathway (Fig. 3o).

### Activation of the AXL-ICD/SIRT2/RUVBL1/2/INO80 axis in astrocytes of the AD patient brain

To examine the clinical relevance of the AXL-ICD/SIRT2/RUVBL1/2/INO80 axis in AD patient brains, we analyzed post-mortem hippocampal tissue from AD patients and age-matched controls by immunohistochemistry and PLA. Immunostaining revealed robust AXL and SIRT2 expression in GFAP-positive astrocytes across hippocampal regions and the adjacent entorhinal cortex, with markedly higher signal intensity in AD samples than in controls (Fig. 4a-d and Supplementary Fig. 9a-e). Notably, both AXL and SIRT2 showed prominent nuclear enrichment in AD astrocytes (Supplementary Fig. 9b, c, and e). Western blot analysis further detected increased levels of the AXL C-terminal fragment (CTF) and ICD in AD tissues, indicating enhanced proteolytic processing of AXL (Fig. 4e, f). PLA analysis revealed significantly increased AXL/SIRT2 interactions in GFAP-positive astrocytes within the AD hippocampus compared with controls, detected in both cytoplasmic and nuclear compartments (Fig. 4g, h), indicating activation of this signaling pathway in AD astrocytes. Consistent with recruitment of chromatin-remodeling machinery, PLA further demonstrated increased nuclear interactions between AXL and RUVBL1 (Fig. 4i, j) and RUVBL2 (Supplementary Fig. 9f, g) as well as AXL and INO80 (Fig. 4k, l) in AD astrocytes. These interactions were largely absent in control tissues, indicating AD-associated engagement of the RUVBL1/2/INO80 chromatin-remodeling complex downstream of AXL signaling. Together, these findings demonstrate activation of the AXL-ICD/SIRT2/RUVBL1/2/INO80 axis in AD astrocytes, supporting its pathological relevance in the human AD brain.

**Fig. 4.**
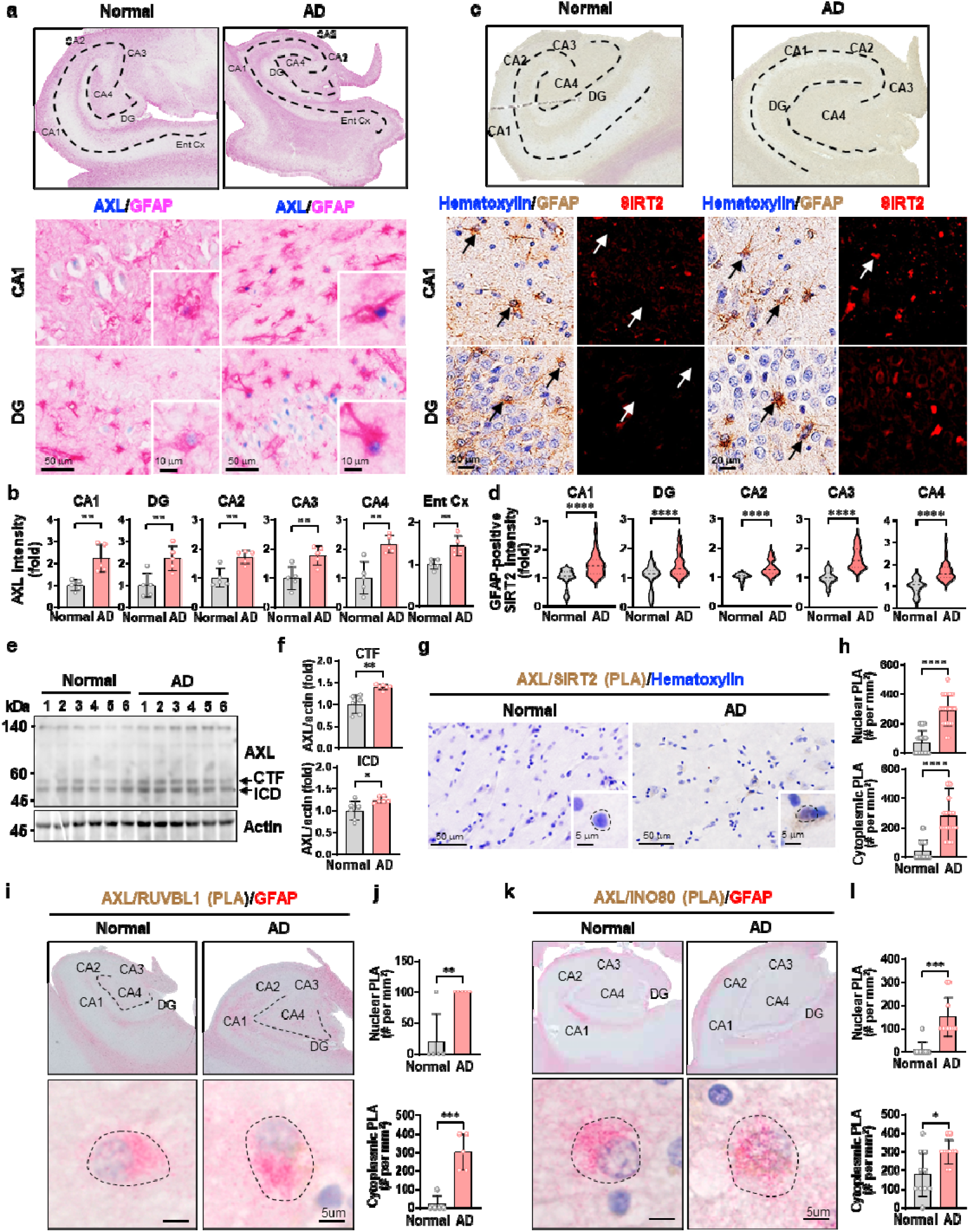
AXL and its interactions are increased in astrocytes of AD brain. **a, b** Immunohistochemical staining of AXL (blue) and GFAP in postmortem hippocampal sections from control subjects and AD patients, with quantification of AXL signal intensity in GFAP-positive astrocytes (*n* = 5 /group). Scale bars, 50 µm (low magnification) and 10 µm (insets) **c, d** Immunohistochemical staining of GFAP and SIRT2 in hippocampal regions of control and AD brains with hematoxylin nuclear counterstaining and quantification of SIRT2 immunoreactivity in GFAP-positive astrocytes (*n* = 40/group). Scale bar, 20 µm. **e, f** Immunoblots of frontal cortex lysates from control and AD brains showing full-length AXL and lower-molecular-weight AXL species corresponding to C-terminal fragment (CTF) and intracellular domain (ICD), with densitometric quantification normalized to actin (*n* = 6/group). **g, h** PLA showing colocalization of AXL and SIRT2 in cortex of control and AD brains and quantification of nuclear and cytoplasmic PLA-positive astrocytes and neurons normalized to tissue area (*n* = 20/group). **i, j** PLA showing colocalization of AXL and RUVBL1 in hippocampal astrocytes of control and AD brains with quantification (*n* = 5/group). **k, l** PLA showing colocalization of AXL and INO80 in hippocampal astrocytes of control and AD brains with quantification (*n* = 10/group). Data are mean ± s.e.m. Statistical significance was determined by unpaired two-tailed Student’s t-test. ns, not significant; *P < 0.05, **P < 0.01, ***P < 0.001, ****P < 0.0001.

### AxSBiP disrupts the AXL-ICD/SIRT2 interaction and restores autophagic flux

Given the functional interaction between AXL-ICD and SIRT2 in autophagy gene induction, we tested whether disrupting this interaction blocks AXL-ICD-dependent transcriptional activation. Structural modeling predicted that the AAAC motif of AXL-ICD directly engages the catalytic domain of SIRT2 (Fig. 5a), providing a structural basis for this interaction, consistent with our mapping data (Fig. 2i, j). Based on the structural modeling (Fig. 2k), we reasoned that competitive binding to the AAAC-binding pocket on SIRT2 would disrupt the AXL-ICD/SIRT2 complex (Fig. 5a). To test this, we designed competitive peptides containing the AAAC motif, extended to a minimum of seven residues to improve specificity, and fused them to a TAT sequence to enable cellular uptake (Supplementary Fig. 10a). Treatment of AXL-ICD-overexpressing U87MG cells with these peptides suppressed AXL-ICD-induced LC3B upregulation (Supplementary Fig. 10b). Notably, the position of TAT conjugation critically influenced inhibitory efficacy, with C-terminal TAT fusion showing greater suppression of LC3B induction than N-terminal fusion, regardless of peptide length (Supplementary Fig. 10a, b). These results indicate that proper exposure of the AAAC motif is required for SIRT2 binding. Competitive targeting of this pocket was therefore sufficient to block AXL-ICD-mediated autophagy. Accordingly, we named the optimized inhibitory peptide as AxSBiP (**Ax**L-ICD/**S**IRT2 **B**inding **i**nhibitory **P**eptide) (Fig. 5b). AxSBiP-TAT potently suppressed AXL-ICD-induced LC3B expression in a dose-dependent manner, with near-complete inhibition at 3 μM (Fig. 5c), and disrupted the AXL-ICD/SIRT2 interaction in co-immunoprecipitation assays (Fig. 5d). Finally, we tested whether AxSBiP-TAT blocks Aβ-induced autophagy protein upregulation and found that it markedly suppressed LC3B and p62 levels (Fig. 5e, f), indicating that the AXL-ICD/SIRT2 complex is required for Aβ-driven autophagy gene activation. Importantly, AxSBiP-TAT did not alter starvation-induced autophagic flux under glucose deprivation, as assessed by bafilomycin A1 treatment (Supplementary Fig. 10c), indicating pathway specificity.

**Fig. 5.**
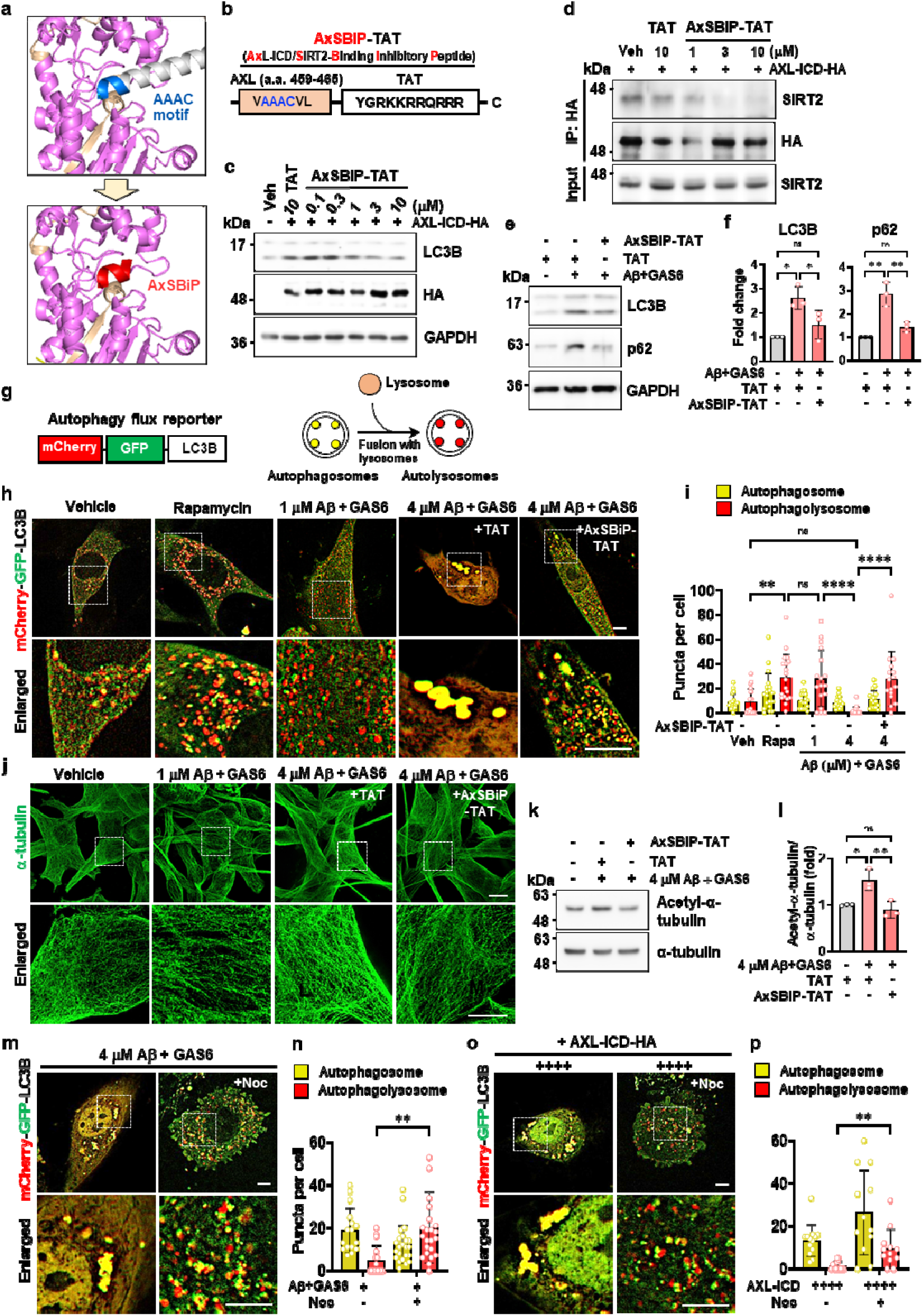
AxSBiP-TAT disrupts the AXL-ICD/SIRT2 interaction and restores autophagic flux by normalizing microtubule acetylation. **a** Docking model of AXL-ICD and SIRT2 interaction showing the AAAC motif at the SIRT2 catalytic domain interface and the predicted binding occupancy of AxSBiP. **b** Schematic of AXL-ICD-derived inhibitory peptides ^35^ containing the AAAC motif (blue), and design of the AxSBiP-TAT peptide, in which TAT serves as a cell-penetrating sequence. **c** Immunoblots of LC3B in AXL-ICD-overexpressing U87MG cells treated with indicated concentrations (0.1–10 µM) of AxSBiP-TAT or control TAT peptide (10 µM). **d** Co-immunoprecipitation of endogenous SIRT2 with HA-tagged AXL-ICD in U87MG cells treated with AxSBiP-TAT (1–10 µM). **e, f** Immunoblots and quantification of LC3B and p62 in U87MG cells treated with Aβ oligomer (4 µM) and GAS6 (250 ng/ml) for 24 h in the presence of TAT or AxSBiP-TAT (5 µM). (*n* = 3). **g** Schematic of the mCherry-GFP-LC3B tandem reporter used to monitor autophagic flux: autophagosomes (GFP⁺/mCherry⁺, yellow) and autolysosomes (mCherry-only, red) **h, i** Restoration of autophagic flux by AxSBiP-TAT. Representative images and quantification of puncta per cell in Aβ-treated U87MG cells. (*n* = 16–19 cells/group). Rapamycin (100 nM) served as a positive control. **j** Immunofluorescence of α-tubulin in Aβ/GAS6-treated U87MG cells with or without AxSBiP-TAT. Scale bars, 10 µm (low magnification) and 5 µm (enlarged). **k, l** Immunoblots and quantification of acetylated α-tubulin (K40) normalized to total α-tubulin in Aβ/GAS6-treated U87MG cells (*n* = 3). **m-q** Representative images (**m, o**) and quantification of puncta per cell (**n, q**) in U87MG cells expressing the autophagy flux reporter. Cells were treated with Aβ/GAS6 (24 h) (**m, n**) or overexpressing AXL-ICD (**o, p**), followed by DMSO or nocodazole (1 µM, 1 h) treatment (*n* = 12–23 cells/group). Scale bars, 10 µm (low magnification) and 5 µm (enlarged). Data are presented as mean ± s.e.m. Statistical significance was determined by one-way ANOVA with Tukey’s post hoc test. ns, not significant; *P < 0.05, **P < 0.01, ****P < 0.0001.

We next examined Aβ-induced autophagic flux using a tandem mCherry-GFP-LC3B reporter that labels early autophagosomes as yellow puncta and autolysosomes as red-only puncta (as GFP fluorescence is quenched in the acidic lysosomal environment) (Fig. 5g). Moderate Aβ exposure (1 μM) increased autophagic flux, as evidenced by accumulation of red-only puncta. This effect was comparable to that of rapamycin, a potent and well-characterized inducer of autophagy that acts by inhibiting the mTOR pathway^35^. In stark contrast, higher Aβ levels (4 μM) reduced autolysosome formation and resulted in enlarged yellow puncta, indicative of a block in autophagic flux at the autophagosome-lysosome fusion step (Fig. 5h, i). Consistently, BafA1-based immunoblot flux assays recapitulated this pattern (Supplementary Fig. 10d). Notably, co-treatment with AxSBiP attenuated the Aβ-induced impairment of autolysosome formation under high-Aβ conditions (Fig. 5h, i). A comparable dose-dependent biphasic pattern was observed with AXL-ICD overexpression, where lower-to-moderate expression progressively enhanced autophagic flux, whereas higher expression impaired it, as confirmed by both BafA1-based immunoblot analysis (Supplementary Fig. 10e) and the tandem LC3 reporter assay (Supplementary Fig. 10f, g). This flux impairment was markedly attenuated by AxSBiP co-treatment (Supplementary Fig. 10f, g). Together, these results indicate that Aβ drives a dose-dependent, biphasic regulation of autophagic flux via the AXL-ICD/SIRT2 axis, promoting autophagy at moderate levels but impairing autophagosome-lysosome fusion beyond a critical threshold.

To determine the mechanism by which AXL-ICD drives biphasic autophagic flux and AxSBiP modulates this response, we focused on AXL-ICD-induced microtubule stabilization, a key determinant of autophagic flux^37^, which occurs via SIRT2 inhibition (Fig. 2n-r). Immunofluorescence analysis showed that while low Aβ or moderate AXL-ICD expression resulted in stabilized and organized microtubules, higher Aβ exposure or elevated AXL-ICD expression led to pronounced microtubule bundling (Fig. 5j and Supplementary Fig. 10h), likely reflecting excessive stabilization. Importantly, AxSBiP partially normalized these structures (Fig. 5j and Supplementary Fig. 10h), a finding further supported by immunoblot analysis showing that AxSBiP reduced the elevated acetylated α-tubulin levels induced by high Aβ or AXL-ICD (Fig. 5k, l and Supplementary Fig. 10i, j). To directly test whether excessive microtubule stabilization is causally responsible for the impaired autophagic flux observed under high-Aβ or high AXL-ICD conditions, we treated cells with nocodazole, a microtubule-destabilizing agent, at doses that partially reverse hyperstabilization without disrupting overall microtubule integrity. Nocodazole treatment restored autophagic flux under both high-Aβ and high AXL-ICD conditions, as evidenced by increased red-only puncta in the tandem LC3 reporter assay (Fig. 5m-p), consistent with a causal link between excessive microtubule stabilization and autophagosome-lysosome fusion defects. Together, these results indicate that AXL-ICD suppresses SIRT2 activity in a dose-dependent manner, leading to increased α-tubulin acetylation and microtubule stabilization; however, under high-Aβ conditions, this stabilization becomes excessive, ultimately disrupting autophagosome-lysosome fusion and impairing late-stage autophagic flux.

### AxSBiP restores autophagic flux, normalizes astrogliosis, and promotes Aβ clearance in APP/PS1 mice

Finally, given that AxSBiP mitigates AXL-ICD-induced microtubule dysregulation, we tested whether this restoration rescues autophagic flux and promotes Aβ clearance. To achieve stable AxSBiP expression *in vivo*, we generated a vector-based construct in which the TAT sequence was replaced with a FLAG tag (Fig. 6a). Lentiviral expression of AxSBiP-FLAG in U87MG cells disrupted the AXL-ICD/SIRT2 interaction (Fig. 6b), indicating that the vector-based construct recapitulates the inhibitory activity of the AxSBiP peptide. Since astrocytic autophagy dysregulation drives H_2_O_2_ overproduction, leading to neuroinflammation and astrogliosis^8,21^, we examined whether AXL-ICD similarly elevates H_2_O_2_ levels and whether AxSBiP can suppress this effect. Using the oROS-G H_2_O_2_ sensor^45^ in AXL-ICD-overexpressing U87MG cells (Fig. 6c), we found that AXL-ICD significantly elevated H_2_O_2_ production, which was completely abolished by AxSBiP, as well as by the MAOB inhibitor KDS2010^46^ and the H_2_O_2_-decomposing hemoglobin enhancer KDS12025^47^ (Fig. 6d), demonstrating that AXL-ICD drives H_2_O_2_ overproduction through the urea cycle-putrescine-MAO-B axis via SIRT2-mediated autophagy.

**Fig. 6.**
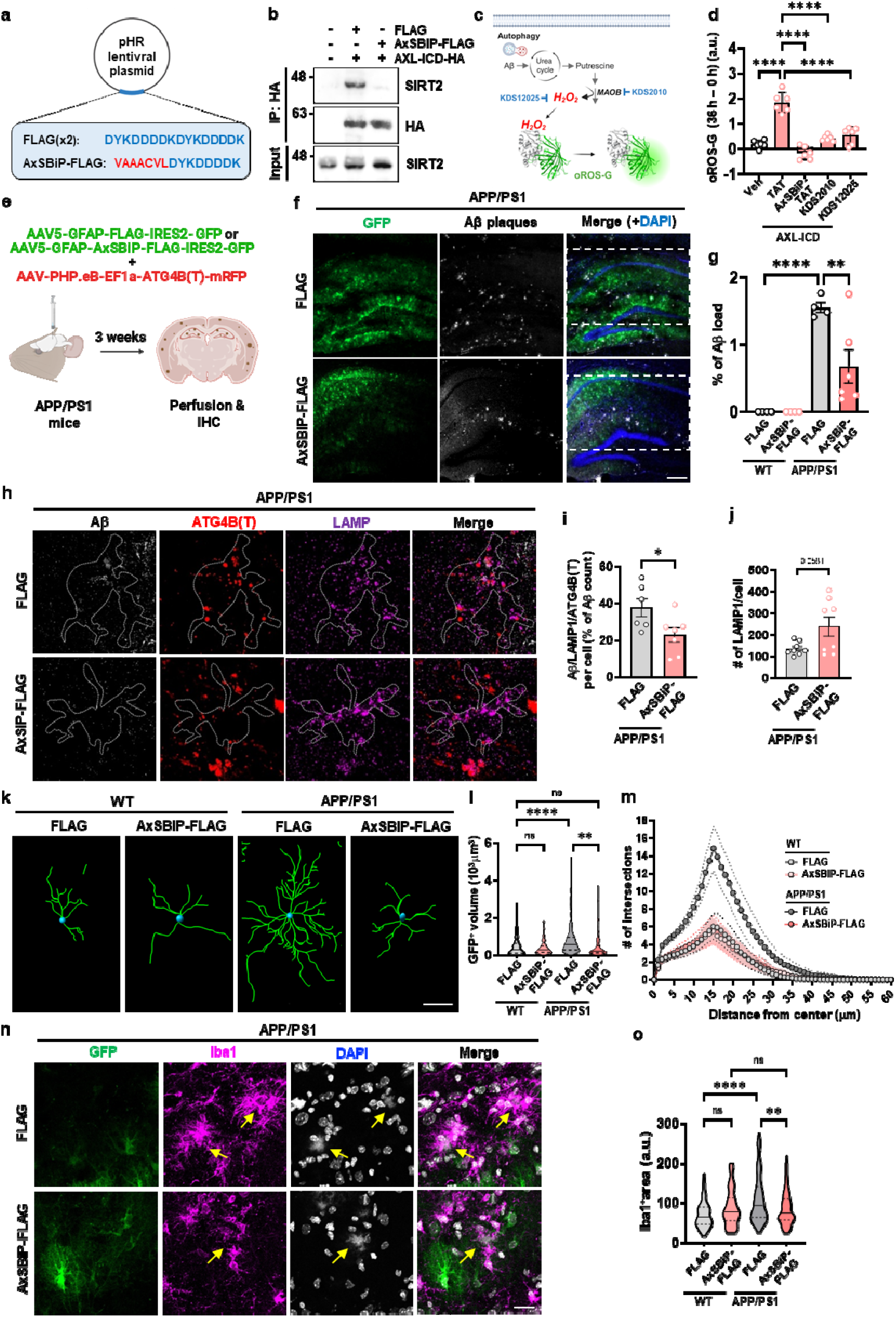
Vectoral AxSBiP expression in astrocytes restores autophagic flux and ameliorates Aβ pathology in APP/PS1 mice. **a** Schematic of the pHR lentiviral plasmid expressing AxSBiP-FLAG. **b** Co-immunoprecipitation of HA-tagged AXL-ICD with SIRT2 and RUVBL1 in U87MG cells co-expressing FLAG or AxSBiP-FLAG. **c** Schematic illustrating autophagy-driven downstream metabolic pathways leading to H_2_O_2_ production in astrocytes. Aβ-induced autophagy activates the urea cycle-linked putrescine metabolic pathway, in which MAOB catalyzes H_2_O_2_ generation. H_2_O_2_ levels were monitored using the oROS-G sensor. The MAOB inhibitor KDS2010 and the H_2_O_2_-decomposing hemoglobin enhancer KDS12025 were tested to validate this pathway. **d** U87MG cells expressing moderate levels of AXL-ICD together with oROS-G were treated with TAT or AxSBiP-TAT (5 µM), or KDS2010 (10 µM) or KDS12025 (1 µM), followed by high-content imaging using an Opera Phenix microscope for 36 h and quantitative analysis. (*n* = 6/group). **e** Schematic of stereotaxic viral delivery into the hippocampus of APP/PS1 mice using AAV5-GFAP-FLAG-IRES2-GFP or AAV5-GFAP-AxSBiP–FLAG-IRES2-GFP together with the autophagy sensor AAV-PHP.eB-EF1α-ATG4B-mRFP. **f, g** Representative images of hippocampal sections from APP/PS1 mice injected with the indicated viruses showing GFP-labelled cells and Aβ plaques, with quantification of Aβ plaque intensity (*N* = 3-4/groups). Scale bar, 200 µm. **h-j** Immunofluorescence of hippocampal sections from APP/PS1 mice showing the autophagy sensor ATG4B and LAMP1 signals in astrocytes (**h**), with quantification of overlapping Aβ, autophagy sensor and LAMP1 signals (*n* = 6-7 cells/group) (**i**) and number of LAMP1 signals per astrocyte (*n* = 8 cells/group) (**j**). **k** Representative reconstruction of GFP-positive astrocytes in the hippocampus using IMARIS. Scale bar, 20μm. **l, m** Truncated violin plot of median and quartiles of the volume of GFP-positive cells and Sholl analysis indicating branching and number of intersections made by GFP-positive cells in the hippocampus of WT and APP/PS1 mice with or without AxSBiP-FLAG expression. (*N* = 2–3/group). Data are presented as mean ± s.e.m. **n, o** Pseudocoloured representative confocal images of mouse hippocampus DG molecular layer stained to visualise AxSBiP/FLAG virus (green), Iba1 (magenta), and DAPI (white). Yellow arrows point to amyloid beta aggregates, also visualised with DAPI. Scale bar, 50 µm (**n**). Truncated violin plot showing median Iba1-positive microglial area in the WT and APP/PS1 animals with AxSBiP or FLAG injection (**o**). Statistical significance was determined by one-way ANOVA with Tukey’s multiple-comparison test (**d**), a two-way ANOVA with Tukey’s post hoc test (**g, l, o**), or unpaired two-tailed Student’s t-test (**i, j**). ns, not significant; *P < 0.05, **P < 0.01, ****P < 0.0001.

We next evaluated whether AxSBiP restores autophagic flux and promotes Aβ clearance *in vivo* using APP/PS1 mice, a well-established AD model with impaired autophagic flux^48,49^. To this end, AxSBiP-FLAG was expressed in astrocytes using AAV5 under the control of the GFAP promoter with an IRES-GFP reporter, and autophagosomes were labeled by co-delivery of an AAV-PHP.eB vector expressing the ATG4B-mRFP sensor^35^ (Fig. 6e). APP/PS1 mice were stereotaxically injected at 50 weeks of age and analyzed three weeks later. In GFAP⁺ astrocytes near Aβ plaques in APP/PS1 mice, AXL was prominently localized within nuclei, whereas it was primarily cytoplasmic in wild-type mice (Supplementary Fig. 11a), mirroring the pattern observed in AD patient brains (Supplementary Fig. 10b, c). Notably, AxSBiP-FLAG expression significantly reduced Aβ plaque burden in GFP-positive regions compared with FLAG controls (Fig. 6f, g), whereas no Aβ plaques were detected in wild-type mice (Supplementary Fig. 11b), demonstrating that disruption of the AXL-ICD/SIRT2 interaction restores Aβ clearance in an AD context. To define the underlying mechanism, we examined whether AxSBiP promotes autophagosome-lysosome fusion in astrocytes of APP/PS1 mice. Consistent with defective autophagic clearance reported in AD^50–52^, FLAG-expressing control mice exhibited accumulation of autophagosome marker-positive puncta together with the lysosomal marker LAMP1 in Aβ-containing astrocytes (Fig. 6h), indicative of impaired autophagic degradation. In contrast, AxSBiP-FLAG expression markedly reduced the proportion of Aβ⁺/autophagosome marker⁺/LAMP1⁺ puncta within GFAP-positive astrocytes (Fig. 6h, i), consistent with enhanced autophagosome-lysosome fusion and subsequent degradation of Aβ-associated autophagic cargo. Additionally, LAMP1 staining was increased in GFAP-positive astrocytes, suggesting an expansion of lysosomal abundance, although this did not reach statistical significance (Fig. 6j). Importantly, AxSBiP-FLAG expression markedly ameliorated astrogliosis in APP/PS1 mice, as evidenced by near-complete restoration of GFP⁺ astrocyte cell volume and branching complexity assessed by Sholl analysis, both of which were substantially elevated in APP/PS1 mice compared to wild-type controls (Fig. 6k-m). This is consistent with our previous findings that astrocytic autophagy dysregulation exacerbates the hypertrophied morphology of astrocytes through either MAOB-dependent^22,8^ or mitophagy defect-induced^8,21^ H_2_O_2_ overproduction, suggesting that Aβ-induced astrogliosis is driven by AXL-ICD through autophagy dysregulation, leading to aberrant H_2_O_2_ overproduction and subsequent neuroinflammation. In addition to astrogliosis, AxSBiP-FLAG expression also reduced microglial activation in APP/PS1 mice, as evidenced by decreased Iba1⁺ area, particularly around Aβ plaques (Fig. 6o, p and Supplementary Fig. 11c). The reduction in microglial activation likely reflects a secondary consequence of the amelioration of astrogliosis, consistent with the bidirectional crosstalk between reactive astrocytes and microglia in neuroinflammation^53,54^. Together, these findings demonstrate that astrocyte-specific AxSBiP expression restores autophagic flux and reduces neuroinflammation, thereby promoting Aβ clearance in an amyloidogenic AD model.

## Discussion

In this study, we demonstrate that Aβ activates a receptor-driven, transcription-dependent autophagy program in astrocytes through regulated intramembrane proteolysis of AXL. Our findings highlight several mechanistic and therapeutic insights (Fig. 7a, b), including 1) AXL as an Aβ-responsive receptor that initiates a *de novo* autophagy program in astrocytes; 2) γ-secretase cleavage generates a nucleus-shuttling, kinase-independent AXL-ICD as a transcriptional effector; 3) previously unrecognized sequence elements within AXL-ICD that coordinate nuclear entry, phase-separated condensate formation, and autophagy gene activation; 4) SIRT2 is repurposed as a nuclear cofactor to drive autophagy gene transcription independently of its deacetylase activity; 5) the AXL-ICD/SIRT2 complex recruits the RUVBL1/2/INO80 chromatin-remodeling machinery to activate autophagy genes; 6) excessive activation of the Aβ-AXL-ICD pathway suppresses SIRT2-dependent control of microtubule remodeling, resulting in α-tubulin hyperacetylation, microtubule hyperstabilization, and functionally halted autophagic flux; and 7) AxSBiP, a competitive inhibitor targeting the AXL-ICD/SIRT2 interface, restores autophagic flux, normalizes astrogliosis, and reduces Aβ plaque burden *in vivo*. Together, these findings define a previously unrecognized signaling axis that links extracellular Aβ sensing to coordinated nuclear activation of autophagy genes and cytoskeletal regulation of autophagic flux in astrocytes. These provide a mechanistic framework for understanding how autophagy transitions from an initially adaptive response to a pathological state during AD progression.

**Fig. 7.**
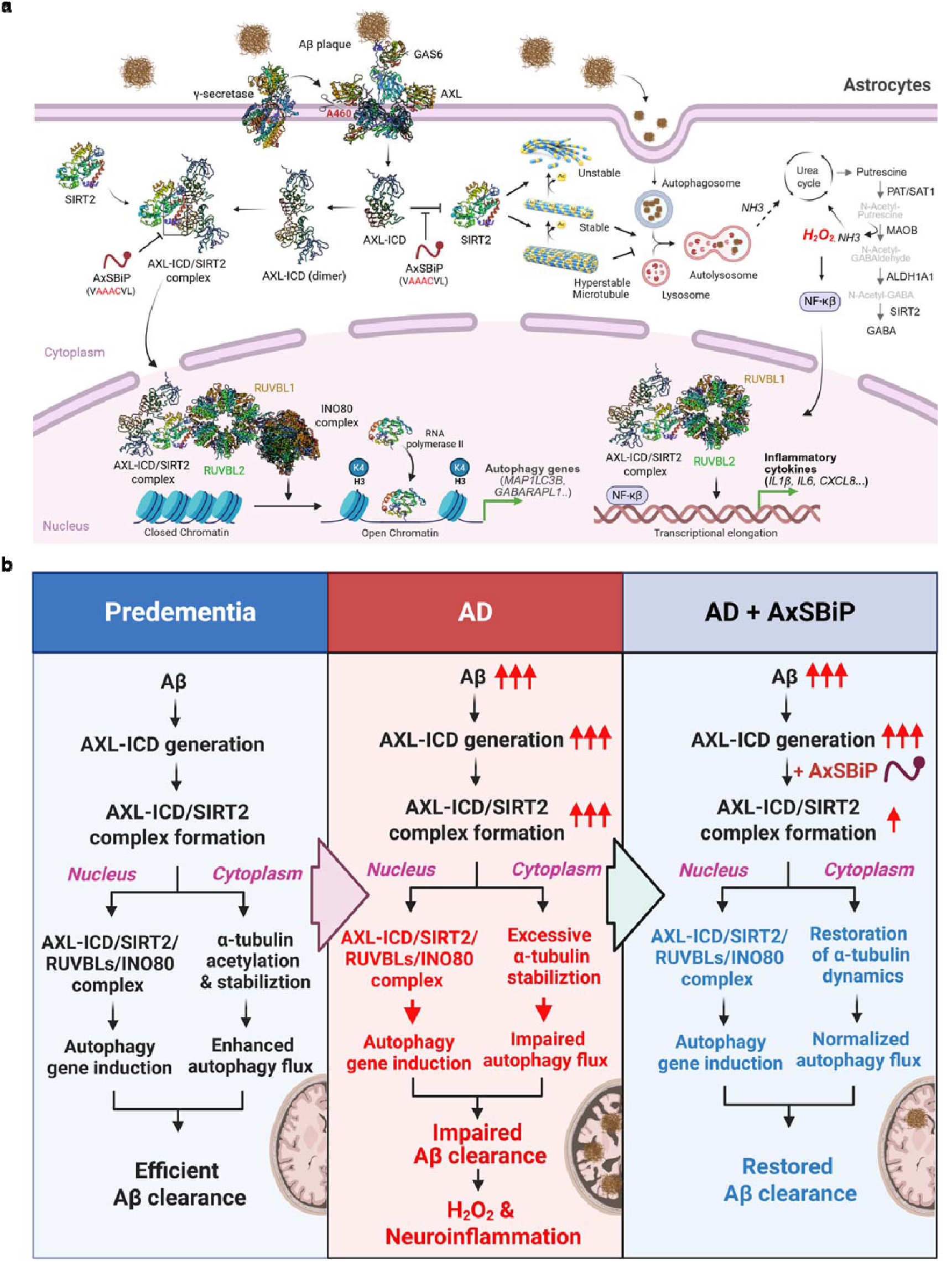
AXL-ICD/SIRT2 signaling links Aβ sensing to astrocytic autophagy and its pathological dysregulation in AD. **a** Proposed mechanism by which Aβ activates a receptor-driven autophagy pathway in astrocytes. Aβ induces γ-secretase-dependent cleavage of AXL to generate AXL-ICD, which forms a complex with SIRT2. In the nucleus, the AXL-ICD/SIRT2 complex recruits the RUVBL1/2/INO80 chromatin-remodeling complex to promote transcription of autophagy genes. In parallel, interaction with SIRT2 inhibits its deacetylase activity, increasing α-tubulin acetylation and stabilizing microtubules to support autophagic flux. Downstream of autophagy, astrocytic urea cycle-linked putrescine metabolism generates H_2_O_2_ and GABA, as previously reported by our group. H_2_O_2_ overproduction and RUVBL1/2 complex activity are proposed to further drive NF-κB-dependent transcription of inflammatory cytokines, including IL1β, IL6, and CXCL8, consistent with prior literature. **b** Pathophysiological model of AXL-ICD signaling during AD progression and its modulation by AxSBiP. In predementia conditions, physiological activation of the AXL-ICD/SIRT2 pathway enhances autophagy and facilitates efficient Aβ clearance. In AD, excessive Aβ signaling elevates AXL-ICD levels, leading to hyperstabilization of microtubules and impaired autophagic flux despite continued autophagy gene induction, resulting in defective Aβ clearance. Defective Aβ clearance under AD conditions is proposed to further drive H_2_O_2_ overproduction and neuroinflammation, consistent with prior literature linking impaired astrocytic autophagy to oxidative stress and inflammatory cascades. Disruption of the AXL-ICD/SIRT2 interaction by AxSBiP restores microtubule dynamics, normalizes autophagic flux and improves Aβ clearance. Illustration created with BioRender.com.

Classical autophagy induction is triggered by nutrient deprivation by signaling pathways such as AMPK activation and mTORC1 inhibition, which regulate the autophagy machinery largely via post-translational modifications^55,56^. In addition, nutrient deprivation can also engage transcriptional programs controlling autophagy genes, including transcription factor TFEB and epigenetic regulators such as CARM1, Tip60, hMOF, G9a and EZH2^42,57–61^. Unlike this canonical pathway, our study identifies a distinct paradigm in which Aβ activates autophagy through a receptor-initiated, transcription-dependent mechanism, independent of nutrient sensing. In this pathway, regulated intramembrane proteolysis converts AXL into a nuclear transcriptional effector that directly drives autophagy gene expression. While canonical AXL signaling drives autophagy through cytoplasmic cascades, such as GAS6-dependent MAPK activation in macrophages^15^ and ROS-AMPK-ULK1 signaling in cancer cells ^14^, our findings reveal a fundamentally distinct mechanism in which Aβ-induced AXL signaling in astrocytes proceeds via regulated intramembrane proteolysis that liberates the AXL-ICD, enabling its nuclear translocation as a transcriptional effector that bypasses conventional kinase signaling. Although TAM receptors have established roles in immune regulation and phagocytosis^62,63^ and have been implicated as Aβ receptors in the brain^11^, their role in autophagy has remained poorly understood. Our results demonstrate that AXL, but not other TAM receptors, is required for Aβ-induced autophagy gene expression and autophagic flux in astrocytes. These findings extend our previous observation that Aβ selectively induces autophagy-related gene expression in astrocytes^8^.

Mechanistically, we show that γ-secretase cleavage converts AXL into a nuclear transcriptional effector. Mapping analyses define the nuclear fragment as AXL-ICD (aa 460– 894) and reveal a compact regulatory architecture composed of an NCS, NES, and NLS that coordinately regulate nuclear localization and exit, condensate assembly, and transcriptional activity. These results demonstrate that nuclear localization alone is insufficient for transcriptional activity; instead, efficient condensate formation via phase separation driven by the AAAC motif is required for autophagy gene activation. Importantly, this motif overlaps with the SIRT2-binding sequence, and SIRT2 recruitment is essential for AXL-ICD condensate assembly. The resulting condensates exhibit features of LLPS-like nuclear compartments, suggesting that SIRT2-dependent condensate formation provides the structural basis for transcriptional activation of autophagy genes. At the transcriptional level, AXL-ICD induces a broad autophagy gene program including *LC3B, p62, GABARAPL1,* and *ATG5* and increases chromatin accessibility at the promoters of a subset of these genes. Notably, this program differs from canonical TFEB-driven pathways, as lysosomal gene expression is not significantly induced despite robust activation of autophagy genes. This uncoupling of autophagy induction from lysosomal biogenesis suggests that AXL-ICD-mediated signaling may increase autophagic input without a corresponding expansion of lysosomal capacity, thereby imposing lysosomal stress under sustained autophagy activation. Accordingly, defining the consequences of this uncoupled autophagy-lysosome response represents an important direction for future investigation.

A key mechanistic insight of our study is that SIRT2 functions as a nuclear cofactor in AXL-ICD-mediated transcription. Although SIRT2 has traditionally been characterized as a cytosolic deacetylase that negatively regulates autophagy^17,64^, our findings reveal a distinct nuclear role. Because SIRT2 lacks a canonical nuclear localization signal ^65,66^, its nuclear entry is likely mediated by interaction with AXL-ICD. Regulated nuclear translocation of SIRT2 has been reported in several contexts, including G2/M progression via CDK1-dependent phosphorylation, bacterial infection, and neurodegenerative signaling^32–34,67^. Here, we demonstrate that AXL-ICD directly binds the catalytic domain of SIRT2 via the AAAC motif, promoting nuclear recruitment and repurposing SIRT2 as a non-enzymatic scaffold for the RUVBL1/2/INO80 chromatin-remodeling complex to activate autophagy genes. In parallel, AXL-ICD binding suppresses the cytosolic deacetylase activity of SIRT2 toward α-tubulin, thereby increasing α-tubulin acetylation and promoting moderate microtubule stabilization to facilitate autophagosome-lysosome fusion^17,36,68,69^. Together, these findings suggest that AXL-ICD bifunctionally reprograms SIRT2 to coordinate nuclear transcription with cytoplasmic autophagy flux. Furthermore, we have identified additional cytoplasmic AXL-ICD interactors, such as ERLIN1/2 and ATAD3A, which are known regulators of autophagosome formation and mitophagy^70–73^. Whether these interactions contribute to AXL-ICD-mediated regulation of autophagosome formation and mitophagy warrants further investigation.

Our present study reveals a paradoxical Janus-like model (Fig. 7b), in which a subthreshold activation of AXL-ICD/SIRT2 axis by Aβ plays a beneficial role in efficient Aβ degradation through autophagy, whereas a suprathreshold activation by overloaded Aβ plays a detrimental role in promoting Aβ accumulation by impairing autophagic flux. Aβ triggers γ-secretase-mediated cleavage of AXL; the resulting AXL-ICD then dually regulates astrocytic responses by transcriptionally activating autophagy genes and modulating microtubule dynamics via SIRT2 inhibition, which reduces α-tubulin deacetylation and consequently increases acetylated α-tubulin levels. This coordinated signaling initially facilitates Aβ degradation. However, upon sustained activation by overloaded Aβ, this AXL-ICD/SIRT2 axis turns pathological by further increasing acetylated α-tubulin levels, rendering microtubules excessively stable and reducing their dynamics. This impairs autophagic flux and prevents Aβ degradation, consequently exacerbating Aβ accumulation. Consistent with this, increased levels of acetylated α-tubulin have been reported in post-mortem AD brain tissues^74^, supporting excessive microtubule stabilization in the AD brain. This model thus explains the paradoxical coexistence of increased autophagosome formation and impaired autophagic flux observed in AD^6,7^. Further supporting this model at the clinical level, soluble AXL (sAXL), the extracellular counterpart generated alongside AXL-ICD during γ-secretase cleavage, shows a stage-dependent pattern: cerebrospinal fluid sAXL correlates with preserved cognitive function in pre-dementia stages, whereas serum sAXL associates with cognitive decline and neurodegeneration in symptomatic AD^75,76^. Collectively, these findings suggest that the AXL-ICD/SIRT2 axis represents a general mechanism by which sustained proteotoxic stress drives maladaptive autophagy despite enhanced autophagosome formation, consistent with the uncoupling between autophagy induction and degradation observed across protein aggregation–associated neurodegenerative diseases^77^.

Within the pathological context, our findings mechanistically extend our prior works linking Aβ-induced autophagy to aberrant astrocytic metabolic rewiring of the urea cycle^23,78^, excessive reactive oxygen species production^22^, and subsequent neurotoxicity in AD^22^ and other related neurodegenerative diseases^47^. Specifically, we now provide evidence that AXL-ICD-mediated autophagy dysregulation drives H_2_O_2_ overproduction in astrocytes. Importantly, H_2_O_2_ overproduction can arise through two distinct mechanisms: MAOB-dependent oxidative stress operating through the urea cycle-putrescine axis downstream of autophagy activation^8,23,78^, and mitophagy defect-induced mitochondrial ROS accumulation^8,23,78^. Together, these findings position AXL-ICD as the upstream molecular trigger that couples extracellular Aβ sensing to H_2_O_2_ overproduction, astrogliosis, and neuroinflammation in AD. Consistent with this, our transcriptomic analysis revealed that AXL-ICD overexpression strongly upregulates NF-κB and a broad set of pro-inflammatory cytokine-related genes (Fig. 1q), which is critical for astrocyte-driven neuroinflammation and neurodegeneration^79,80^. Given that H_2_O_2_ is a well-established activator of NF-κB^81,82^ and that the RUVBL1/2 can further potentiate NF-κB-dependent transcription^83^, AXL-ICD may drive this inflammatory program through H_2_O_2_-mediated NF-κB activation, potentially amplified by RUVBL1/2 (Fig. 7a). Future studies will be required to investigate how the stage-dependent transition from adaptive to maladaptive AXL-ICD activation differentially engages H_2_O_2_-generating mechanisms and how this contributes to the progression of neuroinflammation and neuronal death in AD.

Finally, our findings identify the AXL-ICD/SIRT2 interface as a promising therapeutic target. Analyses of post-mortem human hippocampal tissue demonstrate activation of this signaling axis in astrocytes from AD patients, supporting its clinical relevance. In a mouse model of AD, the competitive inhibitor peptide AxSBiP disrupts the AXL-ICD/SIRT2 interaction, restores the impaired autophagic flux, reduces Aβ plaque burden, and normalizes astrogliosis. This reversal of astrogliosis is likely driven by reduced H_2_O_2_ overproduction, arising from both MAOB-dependent and mitophagy-defect-associated mechanisms, following the restoration of autophagic flux and normalization of autophagy gene expression. Given that H_2_O_2_-driven NF-κB activation and neuroinflammatory cytokine production are key drivers of reactive astrogliosis^84,85^, our findings suggest that AxSBiP attenuates H_2_O_2_-driven neuroinflammation and thereby exerts coordinated beneficial effects on both redox balance and inflammatory signaling.

In summary, growing lines of evidence highlight that reactive astrocytes contribute to AD progression by amplifying neuroinflammation beyond amyloid pathology, supporting the existence of a self-propagating pathological axis driven by reactive astrogliosis and positioning astrocytes as critical and underexplored therapeutic targets in AD^86–88^. In this context, our findings identify the AXL-ICD/SIRT2 axis as a central signaling module linking extracellular Aβ sensing to not only coordinated nuclear and cytoplasmic regulations of the autophagy pathway, but also H_2_O_2_ and cytokine productions leading to neuroinflammation in astrocytes. By demonstrating its maladaptive engagement in both transcriptional hyperactivation of autophagy and SIRT2-dependent microtubule hyperstabilization, accompanied by exacerbated neuroinflammation, we propose that targeting the AXL-ICD/SIRT2 axis with AxSBiP represents a promising therapeutic strategy to reduce both amyloid burden and neuroinflammation. This approach harnesses the endogenous astrocytic autophagic system, promotes Aβ clearance, suppresses H_2_O_2_-driven neuroinflammation, and normalizes reactive astrogliosis, thereby potentially replacing conventional antibody-based Aβ-targeted therapies in AD, without associated neuroinflammation.

## Data availability

The MS-based proteomics data have been deposited in the ProteomeXchange Consortium via the PRIDE partner repository under the dataset identifiers PXD075563 (U87MG) and PXD075596 (human astrocytes). Bulk RNA-seq and ATAC-seq data have been deposited in the Gene Expression Omnibus under accession numbers GSE326144 (RNA-seq) and GSE326145 (ATAC-seq). This paper does not report original code.

## Acknowledgements

We thank Haejin Jung, a senior engineer at the Research Solution Center ^35^, Institute for Basic Science, for performing the flow cytometry analysis. The graphical abstract and all illustrations used in the manuscript were created using BioRender.com. This work was supported by the Institute for Basic Science (Center for Memory and Glioscience, IBS-R001-D2) funded by the Korean Ministry of Science to C.J.L., the National Research Foundation (NRF) Grant from the Korean Ministry of Education, Science and Technology (RS-2022-NR070632 to H.R.) and KIST Grant (26E0122 to H.R.). During the preparation of this work, the author(s) used ChatGPT (OpenAI) and Claude (Anthropic) in order to assist with manuscript editing and language refinement. After using these tools/services, the author(s) reviewed and edited the content as needed and take(s) full responsibility for the content of the published article.

## Authors contributions

T.Y.K. and C.J.L. conceived the project and designed the study. T.Y.K. performed the majority of the experiments, analysed and interpreted the data, and wrote the manuscript with input from all authors. M.B. performed the *in vivo* animal experiments, tissue staining and data analysis. U.P. and S.J.H. prepared human AD brain tissues, performed staining and western blot analyses, and analysed the data under the supervision of H.R. J.L. provided the human AD brain tissues. I.-Y.H. performed bulk RNA-seq and ATAC-seq experiments and bioinformatic analyses. Y.S. and B.L. led the MS-based proteomic analysis. Y.S. performed liquid chromatography and tandem MS data acquisition and analysis. W.Y. performed computational structural modelling of proteins using Schrödinger software. J.-A.L. contributed conceptual input on autophagy flux experiments and provided critical discussion. C.J.L. supervised the study.

## Competing interests

T.Y.K. and C.J.L. have filed a patent application related to the findings described in this manuscript. The remaining authors declare no competing interests.

## Materials and methods

### Reagent

Dimethyl sulfoxide (DMSO) (276855), DAPT (D5942), AGK2 (A8231), 1,6-Hexanediol (240117), bafilomycin A1 (B1793), leptomycin B (L2913), rapamycin (R0395), and nocodazole (M1404) were purchased from Sigma-Aldrich. Amyloid-β (1-42) peptide (Abcam, ab120301), GAS6 (Abnova, H00002621-P01), CB-6644 (BLD Pharm, BD01102615), Ellagic acid (Cayman Chemical, 10569) and EZM2302 (Cayman Chemical, 29954) were obtained from the indicated suppliers. The peptide was purchased from Cusabio (China) and dissolved in sterile H_2_O to prepare a 5 mM stock solution; the peptide sequence is provided in fig. S10A.

### Preparation of oligomeric Aβ42

Amyloid-β (1-42) peptide (Aβ; Abcam, ab120301) was first dissolved in DMSO to prepare a 10 mM stock solution and then diluted to 1 mM in PBS. The peptide solution was incubated at 37 °C for 7 days to allow oligomer formation. Following incubation, samples were aliquoted and stored at −80 °C until use.

### Cell culture

Human astrocytes were obtained from abm (T0280), U87MG cells from the Korean Cell Line Bank (KCLB, 30014) and HEK293T cells from the American Type Culture Collection (ATCC, CRL-3216). Cells were cultured in DMEM (Corning, 10-013-CV) supplemented with 10% FBS (Gibco, 10082147) and penicillin–streptomycin (HyClone, SV30010) at 37 °C in a humidified incubator with 5% CO_2_.

### Plasmids, lentivirus production, and infection

The following plasmids were obtained from Addgene: SIRT1-Flag (#13812), SIRT2-Flag (#13813), SIRT3-Flag (#13814), SIRT4-Flag (#13815), SIRT5-Flag (#13816), SIRT6-Flag (#13817), SIRT7-Flag (#13818), pCDNA-3xFLAG-Pontin (#51635), pcDNA3.1-TFEB-WT-MYC (#99955), pHR-CMV-TetO2_3C-Avi-His6 (#113887), and pHcRed-NPM1wt-C1 (#131818). AXL and SIRT2 deletion constructs and mutant variants were generated using the EZ-Fusion™ HT Cloning Kit (Enzynomics, EZ015TL), whereas point mutations were introduced by site-directed mutagenesis using the EZchange™ Site-directed Mutagenesis Kit (Enzynomics, EZ004S). All constructs were verified by Sanger sequencing (Cosmo Genetech). Primer sequences used for cloning and mutagenesis are available upon request. For gene knockdown, oligonucleotides encoding shRNA sequences targeting the indicated genes, together with the nonspecific control shRNA (all shRNA sequences used in this study, including NS, are listed in Table S1), were cloned into the pLKO.1-puro lentiviral vector. Lentiviral particles were generated by transient co-transfection of shRNA-containing pLKO.1 constructs or expression constructs including pHR-CMV-AXL (FL, full-length; deletion constructs)-HA, pHR-CMV-AXL (FL)-GFP, and pHR-CMV-peptide-FLAG with the packaging plasmid psPAX2 and the envelope plasmid pMD2.G into HEK293T cells using Lipofectamine 3000 (Invitrogen, L3000-015). Viral supernatants were collected at 48 h post-transfection, pooled, and filtered through 0.45-µm pore-size filters. Target cells were incubated with virus-containing supernatants in the presence of polybrene (Merck, TR-1003-G) at a final concentration of 4 µg/ml. After 24 h, the medium was replaced with fresh culture medium, and cells were maintained for an additional 24-48 h before cell lysis.

### Western blot analysis

Proteins were extracted from cells using RIPA lysis buffer (Rockland, mb-030-0050) supplemented with a 1× protease inhibitor cocktail (GenDEPOT, P3100). Lysates were centrifuged at 13,000 rpm for 15 min and the supernatants were analysed by SDS-PAGE. Alternatively, cells were lysed directly in 1x SDS sample buffer and subjected to SDS-PAGE. Separated proteins were transferred onto nitrocellulose membranes (Bio-Rad, 1620115). Membranes were blocked with 5% skim milk (BD, 232100) in Tris-buffered saline with 0.1% Tween-20 (TBST) and incubated overnight at 4 °C with primary antibodies diluted in TBST containing 1% Bovine Serum Albumin (BSA) (GenDEPOT, A0100). The following day, membranes were incubated with HRP-conjugated secondary antibodies for 1 h at room temperature. Immunoreactive bands were visualized using ImageQuant™ LAS 500 (Amersham). Primary antibodies used included: AXL (Cell Signaling, 8661), MERTK (Cell Signaling, 4319) Tyro3 (Cell Signaling, 5585), LC3A (Cell Signaling, 4599), LC3B (Cell Signaling, 3868), ATG5 (Cell Signaling, 12994), SQSTM1/p62 (Santa Cruz, 28359), GABARAP (GeneTex, GTX132657), GABARAPL1 (GeneTex, GTX132664), SIRT2 (Abcam, ab211033), GAPDH (Abcam, ab8245), Lamin B1 (Santa Cruz, 374015), β-Actin (Cell Signaling, 4970), Tip60 (Cell Signaling, 12058), RUVBL1 (Fortis, A304-716A), RUVBL2 (Fortis, A302-536A), INO80 (Proteintech, 18810-1-AP), Erlin1 (ATLAS, HPA011252), Erlin2 (ATLAS, HPA002025), ATAD3A (Abnova, H00055210-D01), WDR5 (Cell Signaling, 13105), CARM1 (Fortis, A300-421A), FLAG (Sigma-Aldrich, M2; 3165), HA (BioLegend, 901501), Acetyl-α-tubulin (Lys40) (Cell Signaling, 3971), α-tubulin (Sigma Aldrich, T6199). All used at a dilution of 1:1000 unless otherwise indicated. Secondary antibodies were HRP-linked anti-rabbit IgG (Seracare, 5450-0010), anti-mouse IgG (5220-0341), anti-rabbit IgG (light chain specific; Jackson ImmunoResearch, 211-032-171) and anti-mouse IgG (light chain specific; 115-035-174).

### Nuclear and Cytosolic Fractionation

Cellular fractionation was performed using the Nuclear/Cytosolic Fractionation Kit (Cell Biolabs, AKR-172) according to the manufacturer’s instructions. U87MG cells expressing mock or HA-tagged AXL-ICD were harvested, washed with ice-cold PBS, and resuspended in ice-cold 1× Cytosol Extraction Buffer containing DTT and protease inhibitors, followed by incubation on ice for 10 min. Cell Lysis Reagent was added, and samples were briefly vortexed and centrifuged at 800 × g for 10 min at 4 °C to obtain the cytosolic fraction. The pellet was washed once and re-centrifuged to minimize cross-contamination. Nuclear proteins were extracted by resuspending the pellet in Nuclear Extraction Buffer containing DTT and protease inhibitors, followed by incubation on ice for 30 min and centrifugation at 14,000 × g for 30 min at 4 °C. The supernatant was collected as the nuclear fraction. Fraction purity was confirmed by immunoblotting using Lamin B1 and GAPDH as nuclear and cytosolic markers, respectively.

### Co-immunoprecipitation (co-IP)

HEK293T or U87MG cells expressing the indicated proteins were generated by lentiviral infection or transient transfection using Lipofectamine 3000 and cultured in 60-mm dishes to ∼80% confluency for 48 h. Cells were washed with PBS, harvested, and lysed in IP lysis buffer (Thermo Fisher, 87788) supplemented with 1× protease inhibitor cocktail and 1× phosphatase inhibitor cocktail (GenDEPOT, P3200). Cell lysates were incubated overnight at 4 °C with 25 μl (bead volume) of anti-HA magnetic beads (Thermo Fisher, 88837). Beads were washed three times with lysis buffer, and bound proteins were eluted in SDS-PAGE sample buffer, boiled, and subjected to SDS-PAGE analysis.

### Quantitative RT-PCR (qRT-PCR)

Total RNA was extracted from cells using TRIzol reagent (Ambion, 15596026) according to the manufacturer’s instructions. Complementary DNA was synthesized from 1 μg of total RNA using SuperiorScript III Reverse Transcriptase (Enzynomics, RT006). Quantitative PCR was performed using Power SYBR Green PCR Master Mix (Applied Biosystems, 4367659) on a QuantStudio™ 2 Real-Time PCR System (Thermo Fisher). Gene expression was normalized to GAPDH, and relative expression levels were calculated using 2^−ΔΔCt method. Primer sequences are provided in Table S2.

### Confocal and lattice structured illumination microscopy

Fluorescence imaging was performed using either confocal or super-resolution microscopy as indicated. For SIRT2 localization analysis (Fig. 2A and C) and Aβ plaque staining (Fig. 6E), images were acquired using an LSM900 confocal microscope (Zeiss) equipped with appropriate laser lines and objectives. All other fluorescence imaging experiments, including AXL-ICD condensates, LC3B puncta, and cytoskeletal structures, PLA puncta, and other immunostaining signals, were performed using a LatticeSIM super-resolution system (Carl Zeiss) equipped with a 63×/1.4 NA oil-immersion objective. Cells were fixed with 4% paraformaldehyde for 15 min at room temperature and permeabilized with 0.1% Triton X-100. After blocking in 2% BSA in PBS, cells were incubated with the indicated primary antibodies overnight at 4 °C, followed by incubation with Alexa Fluor 488- or Alexa Fluor 594-conjugated secondary antibodies (Invitrogen) for 1 h at room temperature. Nuclei were counterstained with DAPI (Thermo Fisher, 62248). Single-plane SIM images were acquired and reconstructed using ZEN software (Zeiss). Quantification of fluorescence intensity or puncta was performed using ImageJ or manual scoring as described in each figure. For live-cell imaging, cells were plated on µ-Dish 35 mm (ibidi, 80136) and imaged using a LatticeSIM super-resolution system equipped with a 63×/1.4 NA oil-immersion objective. Time-lapse images were acquired at 20-s intervals with minimal laser power to reduce photobleaching and phototoxicity.

### Proximity ligation assay ^35^

Protein-protein interactions were examined using the Duolink In Situ Proximity Ligation Assay kit (Sigma-Aldrich, DUO92008) according to the manufacturer’s protocol with minor modifications. U87MG cells expressing mock or AXL-ICD-HA generated using a lentiviral system were cultured on glass coverslips, fixed with 4% paraformaldehyde for 15 min at room temperature, and permeabilized with 0.2% Triton X-100 in PBS for 10 min. After blocking with Duolink blocking solution for 1 h at 37 °C, cells were incubated overnight at 4 °C with primary antibodies, including mouse anti-HA antibody and rabbit antibodies against the indicated proteins. Following washing, cells were incubated with PLA probe PLUS and MINUS secondary antibodies for 1 h at 37 °C. Ligation and rolling-circle amplification reactions were performed using the Duolink ligation and amplification reagents. PLA signals were detected as discrete fluorescent puncta after incubation with the detection reagent. Nuclei were counterstained with DAPI, and samples were mounted using fluorescence mounting medium. Images were acquired using lattice structured illumination microscopy (lattice SIM). PLA puncta were manually counted. For each condition, at least 10 cells from multiple fields were analysed, and the number of PLA puncta within nuclear and cytoplasmic regions was quantified as a measure of protein proximity.

### Cyto-ID autophagy detection assay

Autophagic vesicles were quantified using the Cyto-ID Autophagy Detection Kit (Enzo Life Sciences, ENZ-KIT175) according to the manufacturer’s instructions. U87MG cells were transduced with lentiviral particles expressing INO80-targeting shRNA or non-specific control shRNA for 48 h, followed by transduction with lentiviruses expressing AXL-ICD-HA or control vector (mock) for an additional 48 h, or were directly transduced with AXL-ICD-HA or mock control lentivirus for 48 h without prior shRNA transduction. Cells were harvested, washed with PBS and incubated with the Cyto-ID detection reagent diluted in assay buffer for 30 min at 37 °C protected from light. Cells were then washed once with assay buffer, resuspended in PBS and analysed on a BD LSRFortessa flow cytometer. Data were acquired using the FITC channel and analysed using FlowJo software. Mean fluorescence intensity (MFI) of Cyto-ID staining was used as a quantitative measure of autophagic vesicle accumulation.

### Autophagosome sensor imaging

U87MG cells expressing AXL-ICD or mock control were seeded on glass coverslips and transfected with plasmids expressing the LC3A/B autophagosome sensor (Fyco1-mRFP) or the GABARAP-family autophagosome sensor (ATG4B(Tn)-mRFP) ^31^ using Lipofectamine 3000 according to the manufacturer’s instructions. Twenty-four hours later, cells were fixed with 4% paraformaldehyde for 15 min at room temperature, washed with PBS, and mounted using fluorescence mounting medium containing DAPI for nuclear counterstaining. Images were acquired using a LatticeSIM super-resolution system. Autophagosome sensor-positive puncta were quantified using ImageJ (NIH). Fluorescent puncta with an area larger than 0.5 µm² were counted as puncta, and the number of puncta per cell was calculated.

### oROS-G H_2_O_2_ sensor imaging

U87MG cells were transduced to express the genetically encoded H_2_O_2_ sensor oROS-G^45^ together with AXL-ICD at moderate or high levels. Cells were plated on 96-well polymer-bottom imaging plates (Ibidi, 89626) and treated with TAT or AxSBiP–TAT (5 µM), or with KDS2010 (10 µM) or KDS12025 (1 µM). Live-cell imaging was performed using an Opera Phenix high-content imaging system (PerkinElmer) over 36 h at 20 min intervals, and green fluorescence intensity was quantified and normalized to the number of cells detected in each well. Fluorescence intensity of oROS-G was quantified using Harmony software (PerkinElmer). For analysis, fluorescence values were averaged from the first three time points (early phase) and from three time points at 36 h (late phase) for each cell.

### Autophagy flux assay

U87MG cells were seeded on glass coverslips and transfected with the tandem fluorescent autophagy flux reporter mCherry-GFP-LC3B. Where indicated, cells were treated with TAT or AxSBiP-TAT under the indicated experimental conditions, then fixed with 4% paraformaldehyde for 15 min at room temperature, washed with PBS, and mounted using fluorescence mounting medium. Images were acquired using a LatticeSIM super-resolution system. Autophagic structures were quantified using ImageJ (NIH). GFP-positive/mCherry-positive double-positive puncta were defined as autophagosomes, whereas mCherry-positive/GFP-negative puncta were defined as autolysosomes, based on quenching of GFP fluorescence in acidic lysosomes. Fluorescent puncta with an area larger than 0.5 µm² were included for analysis, and the numbers of autophagosomes and autolysosomes were quantified on a per-cell basis as a measure of autophagic flux.

### RNA- and ATAC-seq library preparation

Total RNA was isolated using the RNeasy Kit (Qiagen) following the manufacturer’s protocol. RNA samples were then used for construction of sequencing libraries. Libraries for RNA-seq were generated with the NEBNext Ultra II RNA Library Prep Kit for Illumina (NEB, E7770) in combination with NEB multiplex oligos and mRNA enrichment using the Dynabeads mRNA DIRECT Kit (Invitrogen, 61012), according to the respective instructions. ATAC-seq libraries were prepared with the ATAC-seq Kit from Active Motif (53150). In brief, freshly harvested cells were lysed in the kit-provided lysis buffer, and 100,000 isolated nuclei were incubated with Tn5 transposase for 30 minutes at 37 °C with shaking at 800 rpm. The transposed fragments were subsequently purified and amplified by PCR using indexed i5/i7 primers, followed by size selection with SPRI beads. Final library quality was evaluated using an Agilent Bioanalyzer, and DNA concentrations were quantified with a Qubit fluorometer (Thermo Fisher Scientific).

### Processing and analysis of bulk RNA-seq data

Initial read quality was assessed with FASTQC (v0.12.1), followed by adaptor and low-quality base removal using Cutadapt (v2.6) with parameters *–q 10 –m 15 –e 0.10*. The filtered reads were aligned to the human reference genome (GRCh38/hg38)) via STAR (v2.7.3a), and gene-level quantification was obtained using the Subread package. Differential expression was evaluated using DESeq2, with transcripts considered significantly regulated at adj. *P < 0.05*. Principal component analysis (PCA) was carried out on variance-stabilized (VST) data generated within DESeq2. Heatmaps were constructed from z-score–scaled normalized counts. For pathway and functional annotation, significantly altered genes were selected based on log2 fold-change and statistical thresholds from the DESeq2 results, and Gene Ontology and pathway enrichment analyses were conducted using the enrichR, and KEGG pathway enrichment was performed using the *enrichKEGG* function from the clusterProfiler package.

### Preprocessing and analysis of ATAC-seq data

Quality assessment of paired-end ATAC-seq reads was carried out using FASTQC (v0.12.1). Adapter and low-quality sequences were removed with Cutadapt (v2.6) using the parameters *–q 10 –m 15 –e 0.10*. Filtered read pairs were aligned to the human reference genome (GRCh38/hg38) using Bowtie2 (v2.2.5) in local, sensitive mode with a maximum fragment length of 2,000 bp. Duplicate fragments were identified with Biobambam (v2.0.87), and alignment files were subsequently processed with SAMtools (v1.6) to remove reads with mapping quality <30, marked duplicates, and improperly paired fragments. Peak calling for each biological replicate was performed in MACS2 using the *BAMPE* format, applying a read shift of –100 bp (*--shift –100*) and an extension size of 200 bp (*--extsize 200*). Consensus peak sets were produced by combining replicate peak files (narrowPeak) and merging overlapping intervals using the GenomicRanges package in R. Annotation of peak locations and genomic distribution analyses were performed with ChIPseeker, and transcription factor motif enrichment was evaluated using FIMO from the MEME Suite.

### Proteomics

Following HA immunoprecipitation (HA-IP) of protein lysates from human astrocytes and U87MG cells, the eluted samples were initially purified and buffer-exchanged into 50 mM Tris-HCl (pH 8.0) using a 3 kDa molecular weight cut-off centrifugal filter. After vacuum drying, protein samples were dissolved in 8 M urea in the same buffer. Subsequently, proteins were reduced with 5 mM Tris(2-carboxyethyl)phosphine hydrochloride (TCEP) and alkylated with 20 mM chloroacetamide (CAA) at room temperature. Trypsin/Lys-C protease mixture (Promega, V5071) was added at a 1:30 enzyme-to-protein ratio. After 4 h at 37 °C, samples were diluted to 1 M urea in the same buffer and digestion was continued overnight. According to the manufacturer’s instructions, tryptic peptides were purified and enriched using C18 solid-phase extraction (Waters, WAT023590). Peptides were loaded onto the cartridge with 0.2 % trifluoroacetic acid (TFA) in water and eluted with 80 % acetonitrile (ACN) in water (v/v), followed by elution with 80 % ACN containing 0.2 % TFA. Vacuum-dried peptide samples were reconstituted in 0.1 % formic acid ^32^ for LC–MS/MS analysis. Each sample was analyzed in triplicate using a nanoLC-MS/MS system (Thermo Fisher Scientific, UltiMate 3000 RSLCnano coupled to an Orbitrap Q Exactive). Tryptic peptides were first trapped on a C18 trapping column (Thermo Fisher Scientific, PN 164535) and subsequently separated on a C18 analytical column (Thermo Fisher Scientific, PN ES903) using an LC gradient with mobile phase A (0.1 % FA in water) and mobile phase B (0.1 % FA in ACN). The gradient conditions were as follows: 5% B (0–15 min), 5–20% B (15–80 min), 20–40% B (80–95 min), and 40–95% B (95–98 min) at a flow rate of 0.3 μL/min. High-energy collision dissociation (HCD-MS/MS) was performed in positive ion mode with an m/z range of 400– 2000, and collision energies were adjusted for precursor ion masses. Raw LC–MS/MS data were processed using MaxQuant and searched against the UniProt human protein database. Search parameters included carbamidomethylation of cysteine as a fixed modification and oxidation of methionine, N-terminal acetylation, and N-terminal carbamylation as variable modifications. Protein identification was based on peptides with a minimum length of six amino acids at an FDR of 0.01, requiring at least one unique peptide per protein group. The Label-Free Quantification (LFQ) algorithm available in MaxQuant was used for relative protein quantification. Gene ontology ^33^enrichment analysis of differentially expressed proteins was performed using the g:Profiler web server.

### Animal husbandry

10-12 months old APP/PS1 animals of both sexes were used in this study, originating from Jackson Laboratory (APPswe/PSEN1dE9; stock number 004462) and maintained as hemizygotes by crossing transgene-carrying mice with B6C3 F1 females. Animals were maintained in12-hr day-night cycle (8am lights on) with food and water provided ad libitum as per the guidelines stated by the Institutional Animal Care and Use Committee of IBS (Daejeon, South Korea). Genotype of the APP/PS1 animals were determined by PCR using the following primers-APP/PS1_F-5’ AAT AGA GAA CGG CAG GAG CA 3’; APP/PS1_R-5’ GCC ATG AGG GCA CTA ATC AT 3’. Age-matched littermates were selected and randomly assigned to the experimental groups.

### Stereotaxic injection

APP/PS1 mice were anesthetized using isoflurane and head-fixed onto stereotaxic frames ^35^. The scalp was incised, and a hole was drilled into the skull above the hippocampus (A/P−1.8, D/V −1.9 from skull surface, M/L ± 1.2 from the bregma). Viruses were combined in a 1:1 ratio and the mixture was loaded into a stainless-steel needle to be injected bilaterally into the dentate gyrus at the rate of 0.1μL/min for 10 min (1μL in each hemisphere). Viruses used: AAV5-GFAP-FLAG-IRES2-GFP, AAV5-GFAP-AxSBiP–FLAG-IRES2-GFP, and AAV-PHP.eB-EF1α-ATG4B-mRFP were generated at the Institute for Basic Science Virus Facility (https://www.ibs.re.kr/virusfacility/). Mice were perfused and brains were excised 2 weeks after injection for immunohistochemistry.

### Mouse brain tissue prep and immunohistochemistry

Animals were anaesthetized using isoflurane and perfused with chilled 0.9% saline, followed by ice-cold 4% paraformaldehyde (PFA) in 0.1 M PBS. The brain was carefully removed and stored in 4% PFA at 4 °C overnight for post-fixation, followed by dehydrolysation in 30% sucrose for 48 hrs. Coronal sections of the hippocampus (30 μm thickness) were prepared in a cryostat and stored in a glycerol-based storage solution at 4 °C till use. Prior to staining, slices were washed in 0.1 M PBS thrice and incubated for 1 hr in blocking solution (4% Donkey Serum, 0.3% Triton X-100 in 0.1 M PBS). Primary antibodies were added to the blocking solution at desired dilution and incubated overnight at 4 °C with gentle rocking, followed by 1 hr incubation at room temperature. Unbound antibodies were washed off by rocking and rinsing the slices with 0.1 M PBS three times, followed by 2 hrs of incubation at room temperature with corresponding fluorescence-tagged secondary antibodies (diluted in blocking solution). Unbound secondary antibodies were washed by rocking and rinsing thrice with 0.1 M PBS, the first wash of which contained 1:1000 DAPI (when required) for nuclear visualization. The slices were then mounted using a fluorescence mounting medium (Dako) and dried. 22–24 μm Z-stacked images in 2 μm steps were processed using the ZEN Digital Imaging for Light Microscopy blue system (Zeiss, ver. 3.2) and ImageJ (NIH, ver. 1.54b) software. Antibodies used in the experiments were as follows (dilutions in blocking solution) - chicken-anti-GFAP (1:500; AB5541, Millipore), rabbit-anti-LAMP1 (1:200, ab211033, abcam), mouse-anti-amyloid beta (1:500; ab126649, abcam), rabbit-anti-AXL (1:200; 8661, Cell Signaling), goat-anti-iba1 (1:500; NB100-1028, Novus), Alexa 647 donkey-anti-chicken anti IgG (1:500, 703–605-155, Jackson), Alexa 488 donkey-anti-rabbit IgG (1:200, 711–547-003, Jackson). Alexa 555 donkey-anti-mouse IgG (1:500, Molecular Probe, A31570); Alexa 647 donkey-anti-mouse IgG (1:500, Jackson, 715-605-150); Alexa 647 donkey-anti-rabbit anti IgG (1:200, Jackson, 711-605-152).

### Human postmortem brain samples

Neuropathological examination of normal and AD postmortem brain samples was conducted according to established protocols of the Boston University Alzheimer’s Disease Research Center (BU ADRC). Neuropathological diagnoses were confirmed and staged using Braak staging criteria. This study was exempted by the Institutional Review Board (IRB) of Boston University School of Medicine (IRB protocol number H-28974), as it involved postmortem tissues not classified as human subjects. All procedures complied with institutional regulatory guidelines and adhered to the principles of the Declaration of Helsinki. Specimen-related information was protected through the BU ADRC system in accordance with NIH policies. Detailed information on the brain tissues is provided in Supplementary Table X.

### Dual chromogenic staining in human postmortem brain sections

First step: Coronal sections (10 µm thick) were prepared from paraffin-embedded postmortem hippocampal tissues obtained from five healthy controls and five patients with AD. Endogenous alkaline phosphatase activity was blocked using BLOXALL® Blocking Solution (SP-6000, Vector Laboratories, USA). Tissue sections were then incubated in 5% bovine serum albumin (A5611, Sigma-Aldrich, USA) for 1 h to prevent nonspecific binding, followed by incubation with anti-AXL antibody (1:200, sc-166269, Santa Cruz Biotechnology, USA) for 24 h. After three washes with PBS, signal amplification was performed using the Vector ABC kit (AK-5000, Vector Laboratories, USA), and AXL immunoreactivity was visualized using Blue substrate (SK-5300, Vector Laboratories, USA).

Second step: Following AXL development, the same sections were incubated with anti-GFAP antibody (1:400, AB5541, Millipore, USA) for 24 h. Detection was performed using Goat Anti-Chicken IgY H&L (Alkaline Phosphatase) (ab6878, abcam, USA) for 2 h at room temperature. GFAP labeling was visualized using Vector Red alkaline phosphatase substrate (SK-5100, Vector Laboratories, USA). The stained tissue sections were sequentially dehydrated through graded ethanol solutions (70%, 80%, 90%, 95%, and 100%), cleared in Histo-clear (HS-200, National Diagnostics, USA), and coverslipped. Dual staining signals, AXL (blue) and GFAP, were examined using a BX63 light microscope (Olympus, Japan) equipped with a DP74 digital camera (1920×1200-pixel; Olympus, Japan).

### In silico modeling

The protein structure of AXL-ICD was generated using AlphaFold2, and the SIRT2 structure was obtained from the Protein Data Bank (PDB ID: 1J8F). Protein structures were prepared for docking simulations at pH 7.4 ± 1 using the Protein Preparation Wizard in Maestro (Schrödinger). Water molecules were removed and hydrogen atoms were minimized using the OPLS4 force field. Protein–protein docking and dimerization simulations were performed using the PIPER module, and the most plausible poses were selected from the top-scoring models. The poses were visualized using the open-source version of PyMOL.

### Statistical analysis

All quantitative data are presented as mean ± s.e.m. Statistical analyses were performed using GraphPad Prism (ver 10.2.3). For comparisons between two groups, unpaired two-tailed Student’s t-test was used. For comparisons among multiple groups, one-way or two-way ANOVA followed by Tukey’s or Sidak’s multiple-comparison test was applied as indicated in the figure legends. The exact statistical test, sample size (n), and number of independent experiments are specified in the corresponding figure legends. Statistical significance was defined as ns, not significant; *P < 0.05, **P < 0.01, ***P < 0.001, and ****P < 0.0001.

## Supplementary Materials

**Figure S1.**
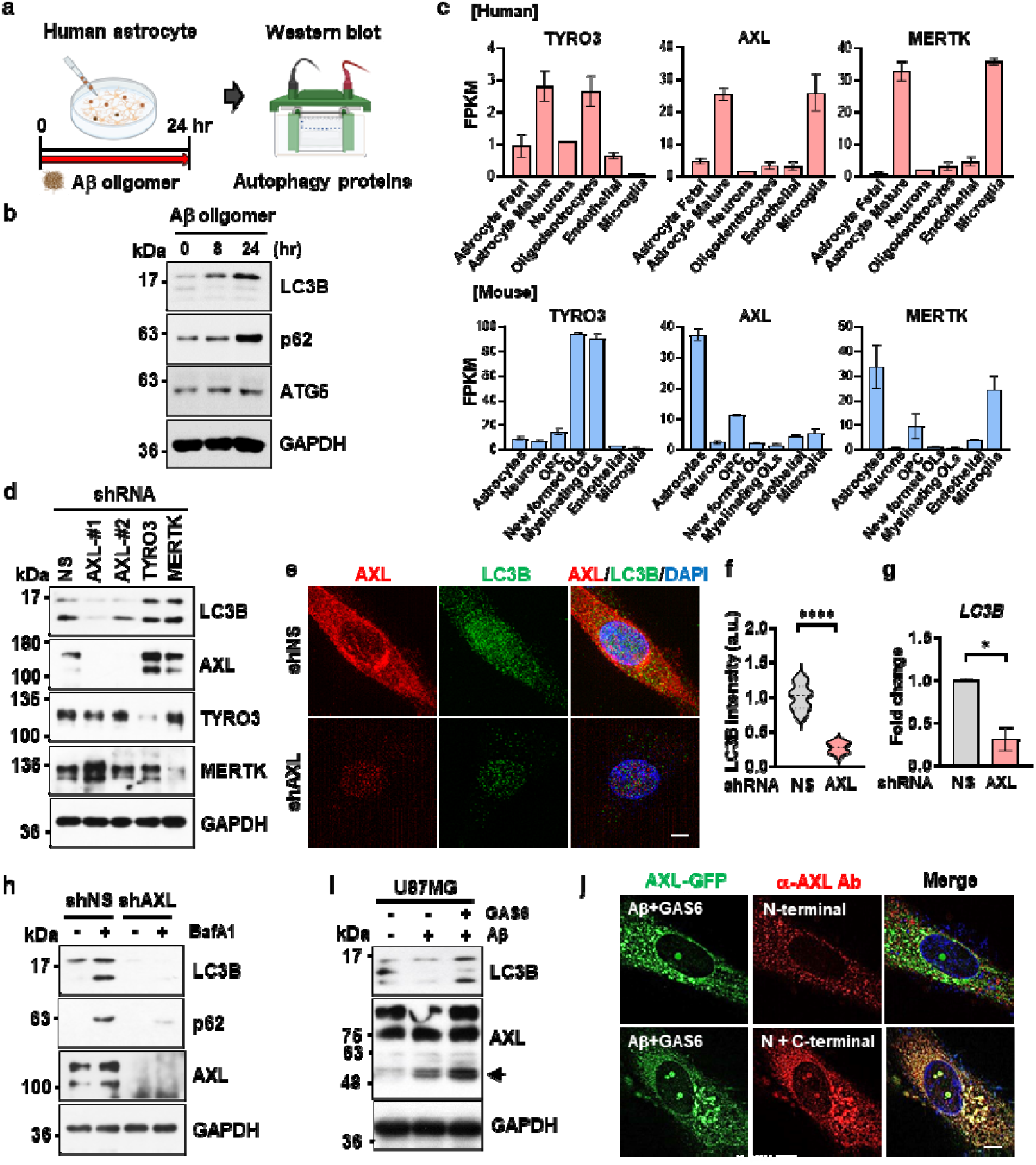
Aβ induces AXL cleavage to generate a nuclear, condensate-forming AXL-ICD that regulates autophagy in astrocytes. **a** Experimental schematic of Aβ oligomer treatment in cultured human astrocytes. **b** Immunoblots of autophagy-related proteins following Aβ oligomers treatment (4 µM, 24 h). **c** Gene expression levels of Tyro3, Axl, and Mertk across major CNS cell populations in mouse and human datasets (https://brainrnaseq.org). OPC and OLs denote oligodendrocyte precursor cells and oligodendrocytes, respectively. Notably, Axl is significantly enriched in astrocytes. **d** Immunoblots of LC3B in human astrocytes following individual knockdown of TAM receptors. GAPDH was used as a loading control. **e, f** Immunofluorescence of AXL and LC3B in human astrocytes expressing shNS or shAXL, with quantification of LC3B signal intensity (*n* = 26-36 cells/group). Scale bar, 5 µm. **g** qRT-PCR analysis of *LC3B* mRNA normalized to GAPDH in shNS- and shAXL-expressing astrocytes (*n* = 2). **h** Autophagic flux analysis in shNS- and shAXL-expressing astrocytes treated with bafilomycin A1 (100 nM, 6 h). **i** Immunoblot of AXL and LC3B in U87MG cells treated with Aβ oligomers (4 μM) with or without GAS6 (250 ng/ml) for 24 h. The arrow indicates AXL-ICD. **j** Immunofluorescence of U87MG cells expressing C-terminal GFP-tagged full length of AXL treated with Aβ oligomers (4 μM) and GAS6 (250 ng/ml) for 24 h stained with antibodies recognizing the N-terminal region or both N- and C-terminal regions of AXL. Scale bar, 5 µm. Data are mean ± s.e.m. Statistical significance was determined by unpaired two-tailed Student’s t-test. *P < 0.05, ****P < 0.0001.

**Figure S2.**
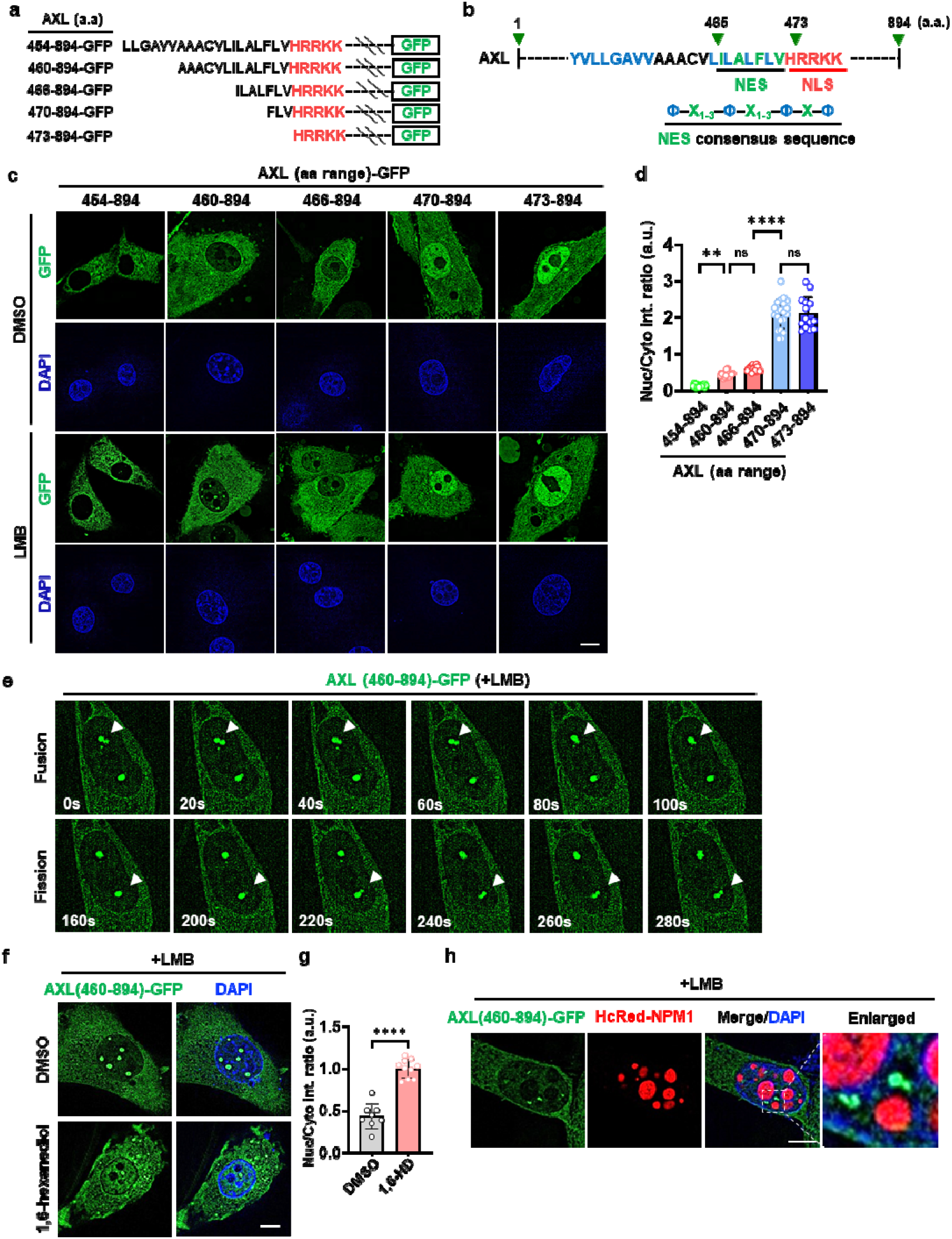
AXL-ICD forms dynamic, liquid-like nuclear condensates. **a** Schematic representation of AXL deletion constructs fused to C-terminal GFP. **b** Schematic of the AXL intracellular domain showing a putative nuclear export signal (NES) and nuclear localization signal (NLS). The predicted NES conforms to the classical consensus motif, where Φ represents hydrophobic residues (L, I, V, F or M). **c, d** Fluorescence of U87MG cells expressing C-terminal GFP-tagged AXL deletion constructs treated with DMSO or leptomycin B (LMB, 10 μM, 16 h) (C). Quantification of nuclear-to-cytosolic GFP intensity ratios, measured from defined nuclear regions and size-matched cytoplasmic regions in U87MG cells expressing each AXL deletion construct (*n* = 13-20 cells/group) **e** Time-lapse live-cell imaging of AXL-ICD-GFP nuclear condensates in U87MG cells treated with LMB (10 μM, 16h). Arrowheads indicate representative condensates undergoing fusion and separation over time, consistent with liquid-like condensate dynamics. Time is indicated in seconds. **f, g** Representative fluorescence images of nuclear condensates in U87MG cells expressing AXL-ICD-GFP and treated with LMB (10 µM, 16 h), followed by treatment with DMSO or 1,6-hexanediol (1,6-HD; 5%, 10 min). Nuclei are stained with DAPI. Scale bar, 5 µm, with quantification of GFP intensity ratios between the condensate-excluded nuclear region and an equal-area cytoplasmic region before and after 1,6-HD treatment (*n* = 8-10 cells/group). **h** Fluorescence images of nuclear condensates in U87MG cells co-expressing AXL-ICD-GFP and HcRed-NPM1 and treated with LMB (10 µM, 16 h). Scale bar, 5 µm. Data are mean ± s.e.m. Statistical significance was determined by one-way ANOVA with Tukey’s multiple-comparison test (**d**) and unpaired two-tailed Student’s t-test (**g**). ns, not significant; **P < 0.01, ****P < 0.0001.

**Figure S3.**
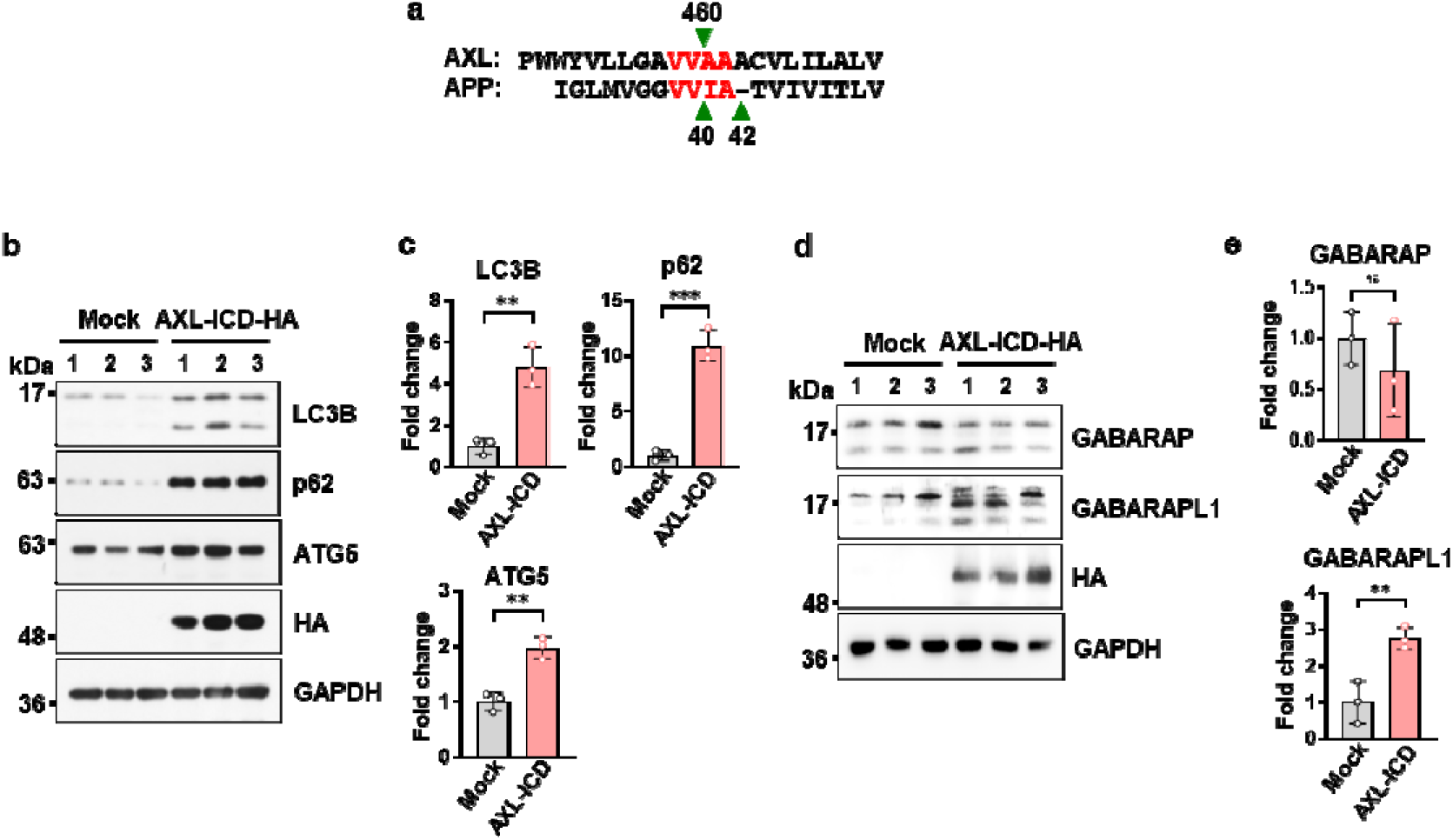
AXL intracellular domain shares γ-secretase cleavage homology with APP and induces autophagy regulation. **a** Sequence alignment showing similarity between the γ-secretase cleavage region of AXL and that of amyloid precursor protein **b-e** Immunoblot analysis of autophagy-related proteins in U87MG cells overexpressing mock or HA-tagged AXL-ICD (**b, d**), with quantification normalized to GAPDH (*n* = 3) (**c, e**).

**Figure S4.**
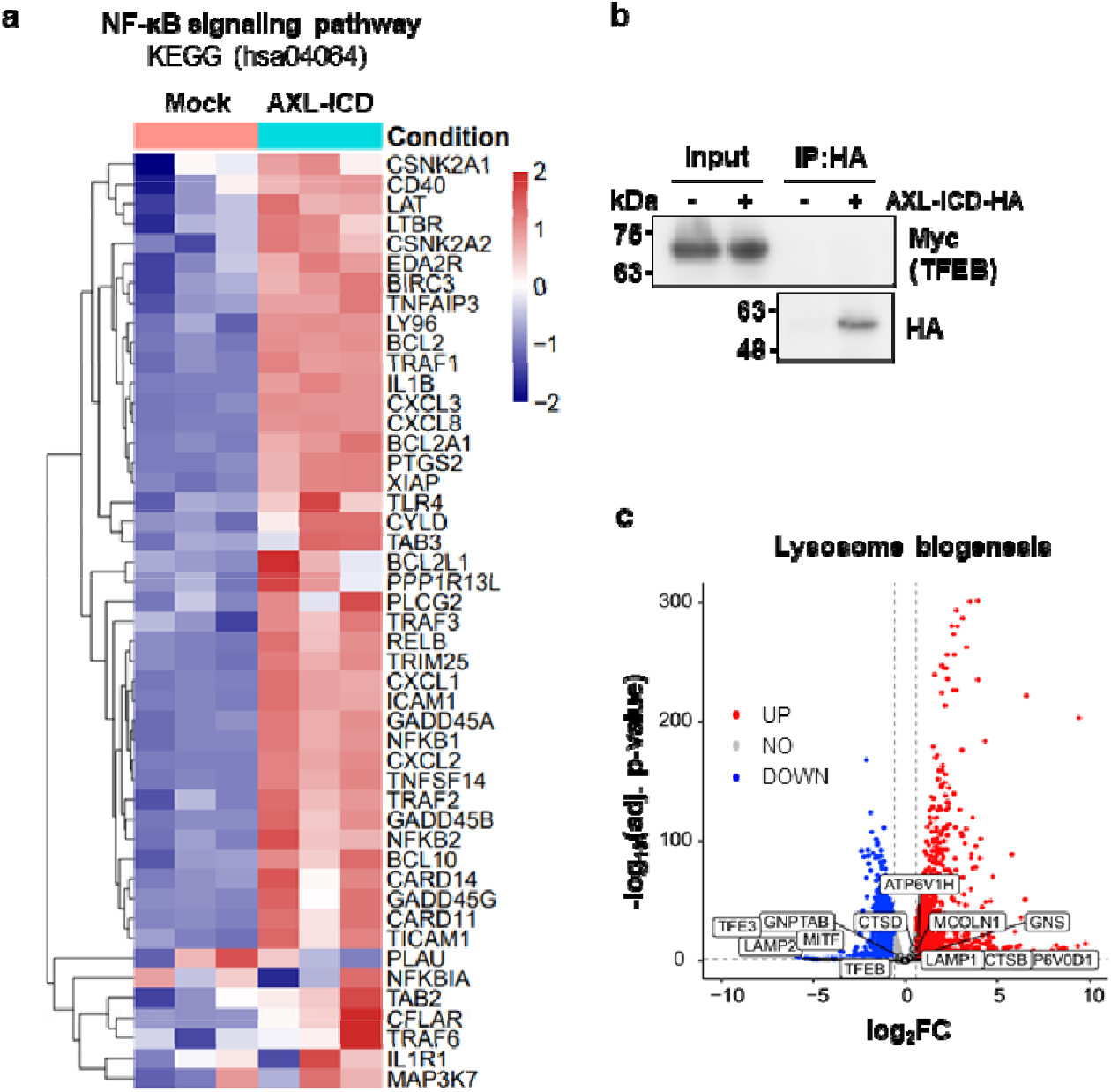
AXL-ICD promotes NF-κB signaling but not TFEB-associated lysosomal programs. **a** Heatmap of NF-κB signaling pathway genes (KEGG: hsa04064) using FPKM-normalized expression values in mock- and AXL-ICD-overexpressing U87MG cells. **b** Co-immunoprecipitation of Myc-tagged TFEB with HA-tagged AXL-ICD in HEK293T cells. **c** Volcano plot showing differential expression of TFEB-associated lysosomal genes in AXL-ICD-overexpressing U87MG cells. Red and blue dots indicate significantly upregulated and downregulated genes, respectively (fold change > 2, adjusted P < 0.05).

**Figure S5.**
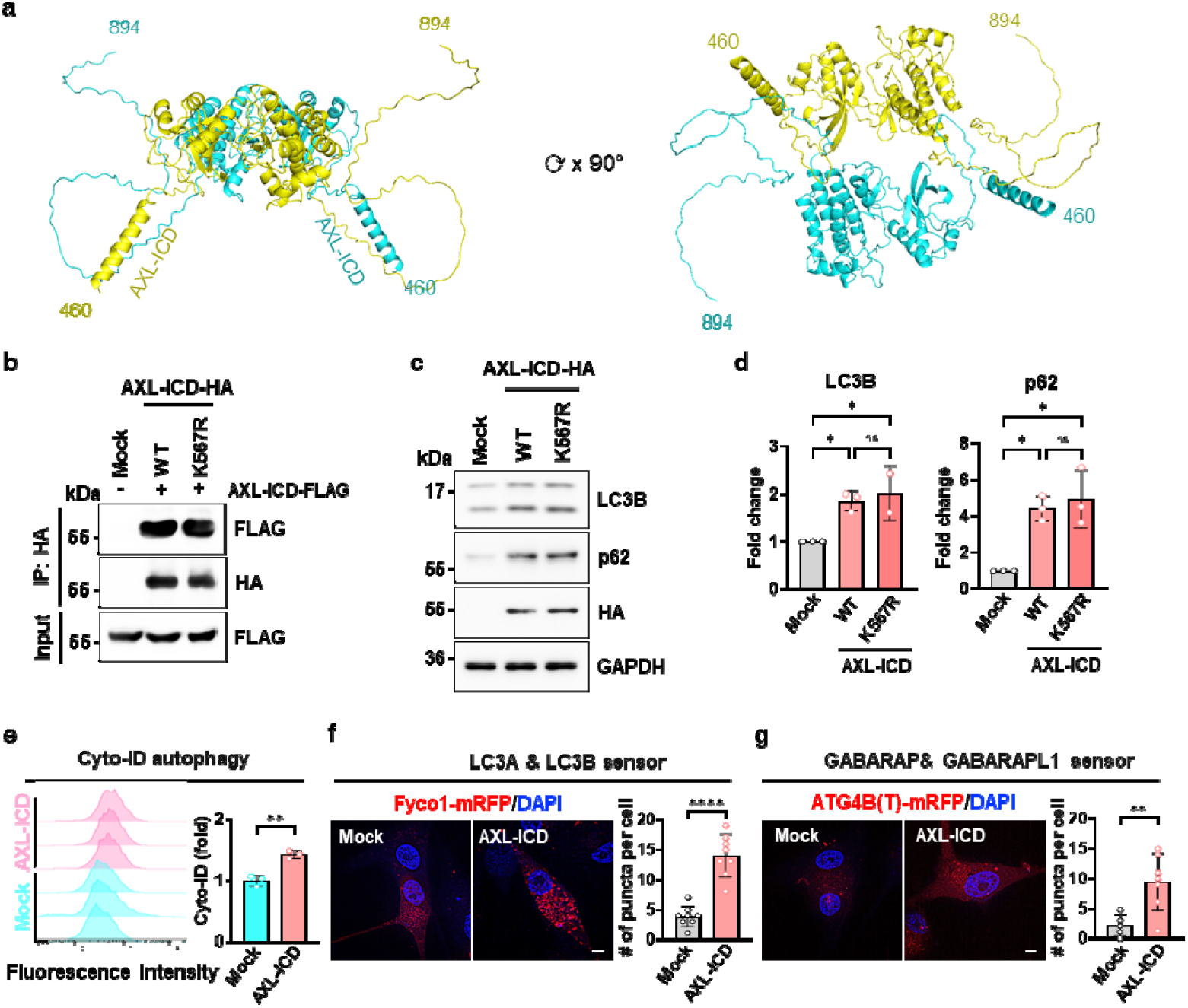
AXL-ICD forms a dimer and promotes autophagosome formation in a kinase-independent manner. **a** Predicted dimeric structure of AXL-ICD (aa 460-894) generated using the Schrödinger modeling suite and visualized in PyMOL. **b** Co-immunoprecipitation showing dimerization of HA-tagged AXL-ICD WT or kinase-dead K567R with FLAG-tagged AXL-ICD in HEK293T cells. **c, d** Immunoblot analysis of LC3B and p62 in U87MG cells expressing WT or kinase-dead K567R AXL-ICD, with quantification (*n* = 3). **e** Flow cytometry of Cyto-ID autophagy signal in mock- and AXL-ICD-overexpressing U87MG cells with quantification of mean fluorescence intensity (MFI) (*n* = 3). **f, g** Fluorescence of LC3A/B (Fyco1-mRFP) and GABARAP-family (ATG4B(Tn)-mRFP)-based autophagosome sensors in mock- and AXL-ICD-expressing U87MG cells, with quantification of mRFP-positive puncta per cell (*n* = 5-8 cells/group). Scale bar, 5 µm. Data are mean ± s.e.m. Statistical significance was determined by one-way ANOVA with Tukey’s multiple-comparison test (**d**) and unpaired two-tailed Student’s t-test ^39^. ns, not significant; *P < 0.05, **P < 0.01, ***P < 0.001.

**Figure S6.**
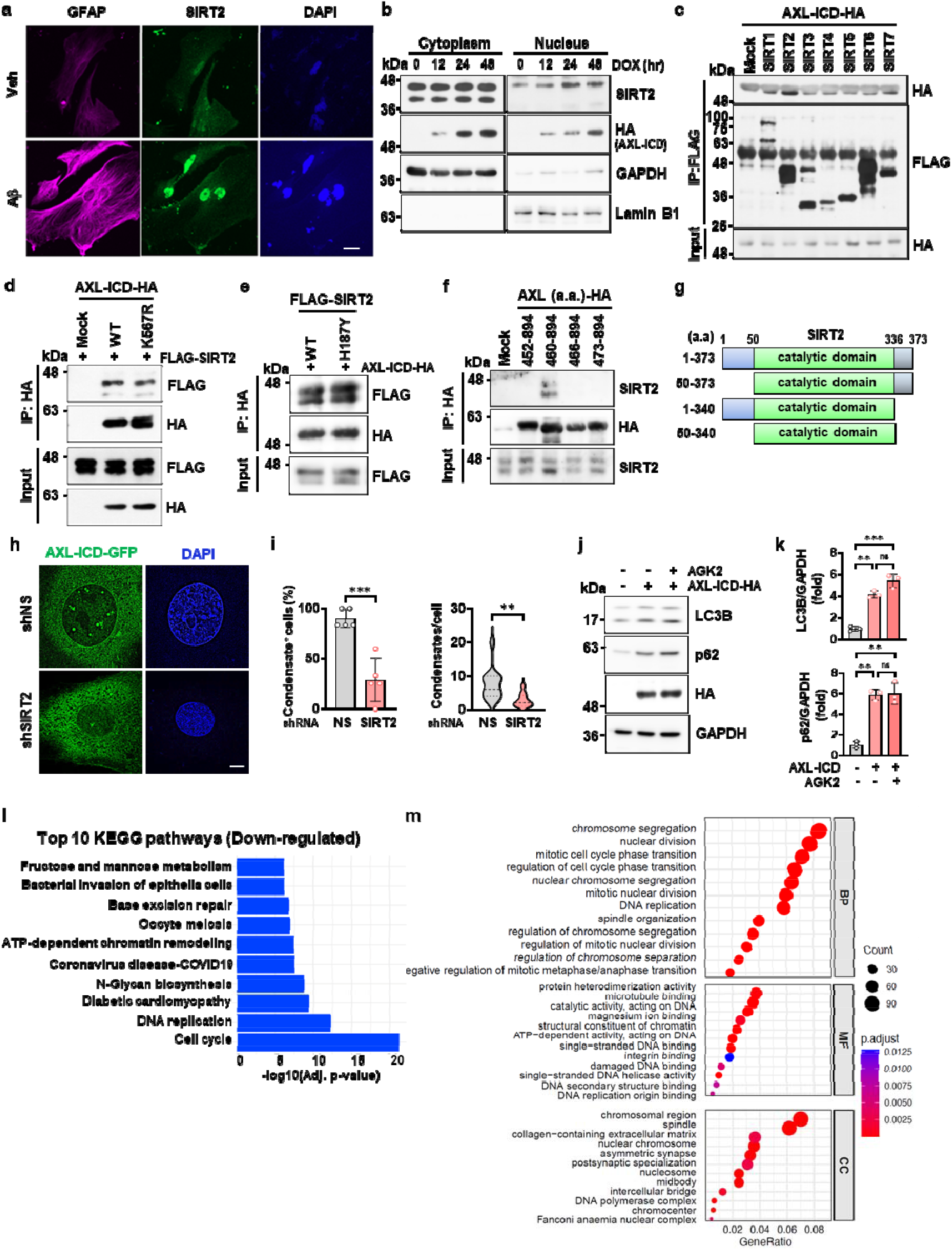
AXL-ICD interacts with SIRT2 independently of enzymatic activity, forms SIRT2-dependent nuclear condensates, and suppresses SIRT2 deacetylase function. **a** Immunofluorescence of primary cortical astrocytes treated with Aβ oligomers (1 µM, 24 h) and stained for GFAP and SIRT2. Scale bar, 5 µm. **b** Immunoblots of SIRT2 in nuclear and cytoplasmic fractions of AXL-ICD-overexpressing U87MG cells. **c** Co-immunoprecipitation of FLAG-tagged sirtuin proteins with HA-tagged AXL-ICD in HEK293T cells. **d, e** Co-immunoprecipitation of HA-tagged AXL-ICD (WT or kinase-dead K567R) with FLAG-tagged SIRT2 (WT or catalytic mutant H187Y) in HEK293T cells. **f** Co-immunoprecipitation of endogenous SIRT2 with HA-tagged AXL-ICD deletion constructs in U87MG cells. **g** Schematic of SIRT2 constructs including full-length and deletion mutants. **h, i** Fluorescence of AXL-ICD-GFP nuclear condensates in NS- or SIRT2-knockdown U87MG cells treated with LMB (10 µM, 24 h), with quantification of the percentage of nuclear condensate-positive cells and the number of condensates per cell (*n* = 5-25 cells/ group). **j, k** Immunoblots of LC3B and p62 in mock- or AXL-ICD-overexpressing U87MG cells treated with AGK2 (5 µM, 24 h), with quantification (*n* = 3). **l** KEGG pathway enrichment analysis of genes downregulated upon AXL-ICD overexpression in U87MG cells, showing enrichment of cell cycle and DNA replication pathways. **m** Gene ontology enrichment analysis of downregulated genes across Biological Process (BP), Molecular Function (MF), and Cellular Component categories. Data are mean ± s.e.m. Statistical significance was determined by unpaired two-tailed Student’s t-test (**i**) and one-way ANOVA with Tukey’s multiple-comparison test (**l**). ns, not significant; **P < 0.01, ***P < 0.001.

**Figure S7.**
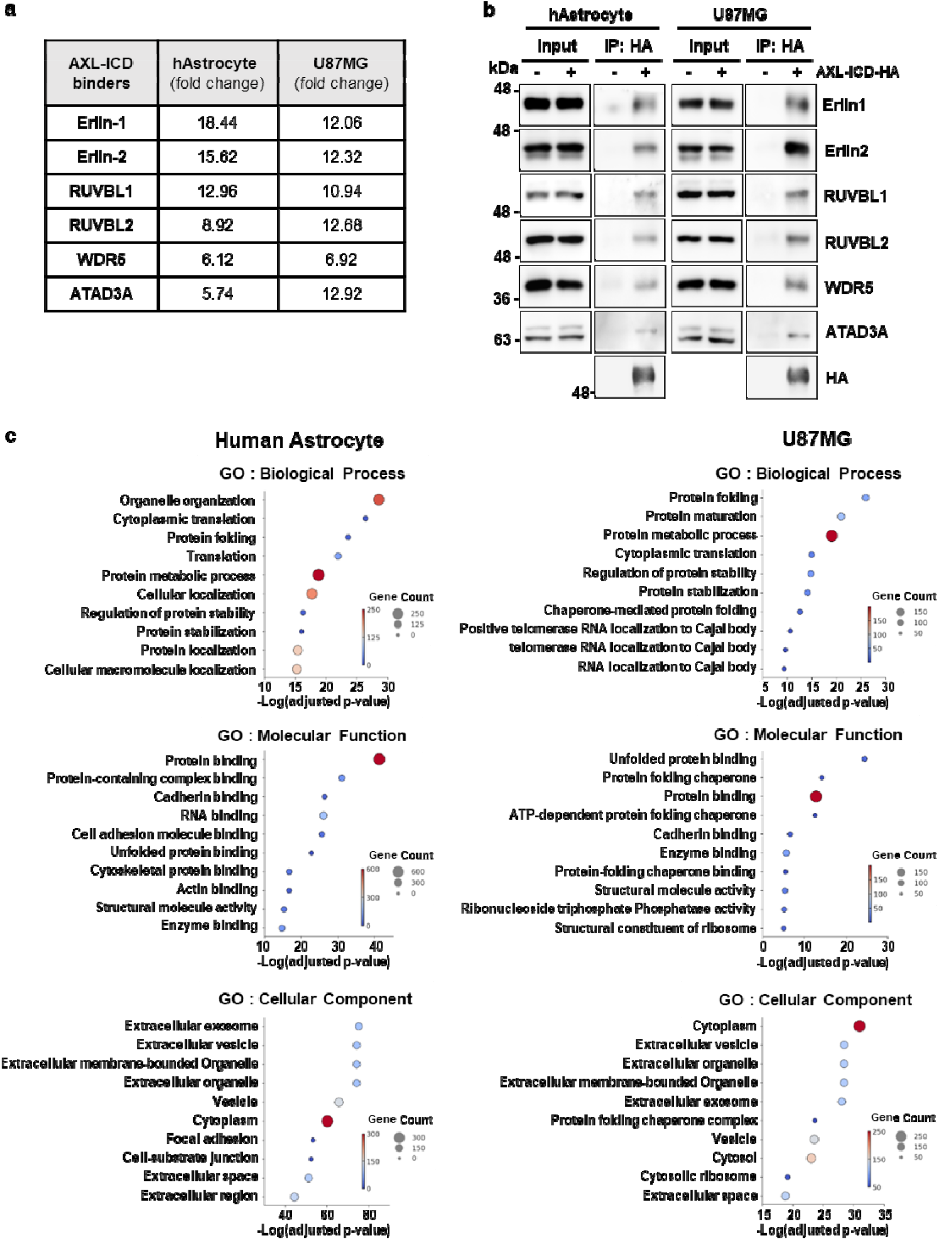
Functional annotation and validation of AXL-ICD-interacting proteins. **a** Proteomic profiling of AXL-ICD-interacting proteins showing fold changes of shared and cell-type-specific interactors in human astrocytes and U87MG cells. **b** Validation of selected AXL-ICD interactors by co-immunoprecipitation and immunoblotting in human astrocytes and U87MG cells. **c** Gene ontology (GO) enrichment analysis of AXL-ICD-interacting proteins identified in human astrocytes (left) and U87MG cells (right), categorized by Biological Process, Molecular Function, and Cellular Component. Dot size indicates gene count; color represents enrichment score.

**Figure S8.**
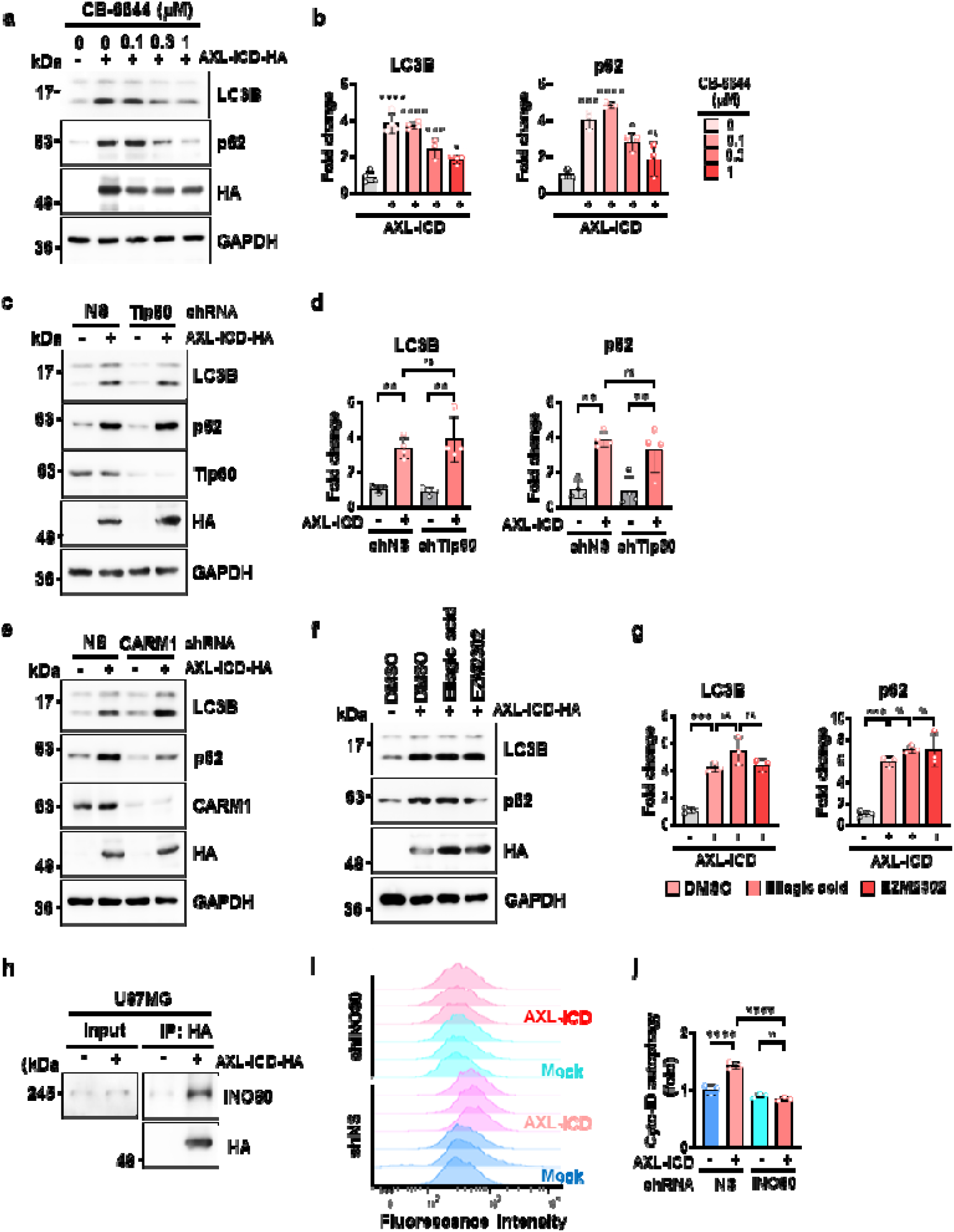
AXL-ICD-driven autophagy induction depends on the RUVBL1/2/INO80 complex. **a, b** Immunoblots of LC3B and p62 in U87MG cells expressing mock or HA-tagged AXL-ICD treated with increasing concentrations of the RUVBL1/2 inhibitor, CB6644, with quantification of relative levels normalized to mock (*n* = 3-4). **c, d** Immunoblots of LC3B and p62 in U87MG cells overexpressing mock or HA-tagged AXL-ICD under NS or Tip60-knockdown conditions, with quantification (*n* = 4). **e-g** Immunoblots of LC3B and p62 in U87MG cells expressing mock or HA-tagged AXL-ICD under NS or CARM1 knockdown conditions (**e**) or following treatment with CARM1 inhibitors (ellagic acid or EZM2302; 10 µM, 24 h) (**f**), with quantification of relative levels normalized to mock for (F) (*n* = 3) (**g**). **h** Co-immunoprecipitation of HA-tagged AXL-ICD with endogenous INO80 in U87MG cells. **I, j** Flow cytometry analysis of Cyto-ID autophagy signal in U87MG cells expressing NS or INO80 shRNA with or without AXL-ICD, with quantification of MFI (*n* =3). Data are mean ± s.e.m. Statistical significance was determined by one-way ANOVA with Dunnett’s multiple-comparison test (**b, g**) and two-way ANOVA with Tukey’s multiple-comparison test (**d, j**) and ns, not significant; **P < 0.01, ***P < 0.001, ****P < 0.0001.

**Figure S9.**
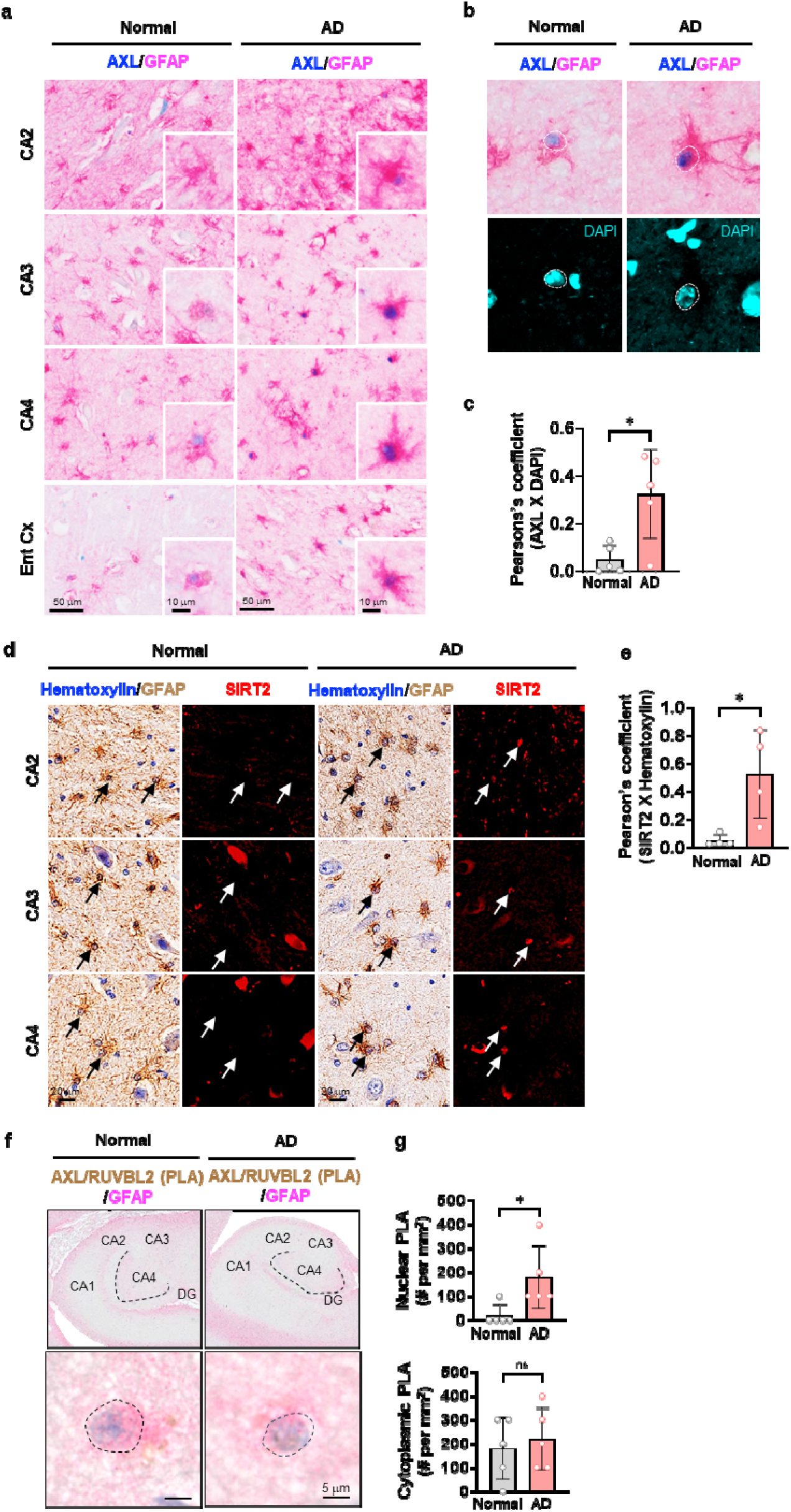
Nuclear enrichment of AXL, SIRT2 and AXL-RUVBL2 interactions in astrocytes in AD. **a** Immunohistochemical staining of AXL and GFAP in hippocampal regions and entorhinal cortex from control and AD postmortem brains. **b, c** Higher-magnification images showing colocalization of AXL with DAPI in GFAP-positive astrocytes with quantification of Pearson correlation coefficients (*n* = 5/group). **d, e** Immunohistochemical staining of GFAP and SIRT2 with hematoxylin nuclear counterstaining in hippocampal regions of control and AD brains with quantification of SIRT2 nuclear localization (*n* = 4/group). **f, g** PLA showing colocalization of AXL with RUVBL2 in hippocampal astrocytes from control and AD brains with quantification of PLA-positive cells normalized to tissue area (*n* = 5/group). Data are mean ± s.e.m. Statistical significance was determined by unpaired two-tailed Student’s t-test. ns, not significant; *P < 0.05.

**Figure S10.**
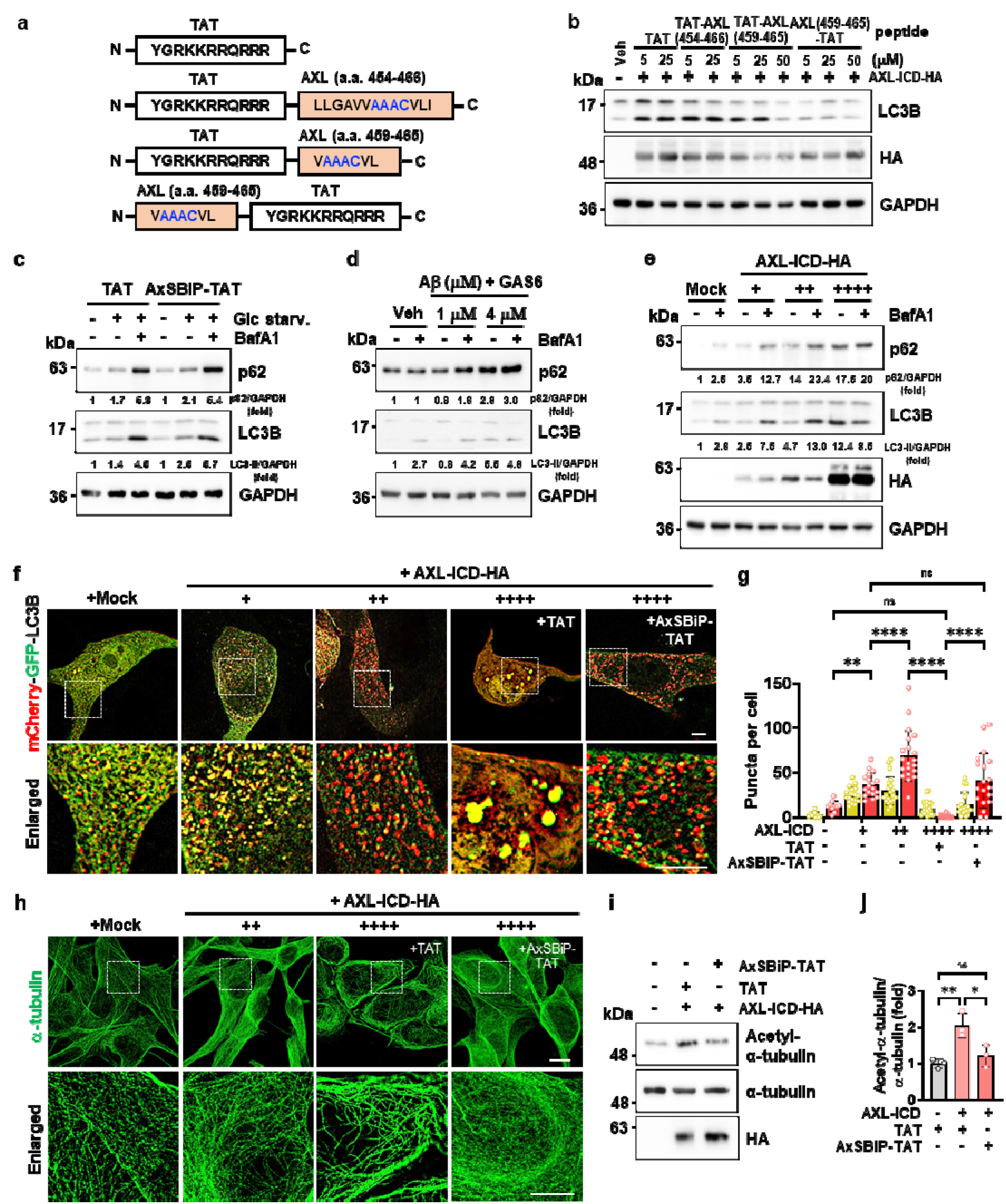
Excess Aβ or AXL-ICD impairs autophagic flux through microtubule hyperstabilization. **a** Schematic of AXL-ICD-derived inhibitory peptides ^35^ containing the AAAC motif (blue), and design of the AxSBiP-TAT peptide, in which TAT serves as a cell-penetrating sequence. **b** Immunoblots of LC3B in U87MG cells overexpressing AXL-ICD-HA and treated with the indicated inhibitory peptides for 24 h. HA indicates AXL-ICD. **c** Immunoblots of LC3B and p62 in U87MG cells treated with TAT or AxSBiP-TAT (5 µM, 18 h) during glucose starvation and subsequently with bafilomycin A1 (100 nM, 6 h). **d, e** Immunoblots of LC3B and p62 in U87MG cells treated with Aβ (1 or 4 µM) and GAS6 (250 ng/ml) for 24 h or expressing increased levels of AXL-ICD, followed by bafilomycin A1 (100 nM, 6 h). **f, g** U87 cells expressing the autophagy flux reporter and overexpressing increasing levels of AXL-ICD were treated with TAT or AxSBiP-TAT (5 µM) for 24 h. Scale bars, 10 µm (low magnification) and 5 µm (enlarged), with quantification of yellow (autophagosomes) and red-only (autolysosomes) puncta per cell (*n* = 10-26 cells/ group). **h** Immunofluorescence of α-tubulin in U87MG cells overexpressing moderate or high levels of AXL-ICD in the presence of TAT or AxSBiP-TAT (5 µM). Representative images and enlarged regions are shown. Scale bars, 10 µm (low magnification) and 5 µm (enlarged). i, j Immunoblots of acetylated α-tubulin (K40) in U87MG cells expressing high levels of AXL-ICD and treated with TAT or AxSBiP-TAT (5 µM), with quantification of acetylated α-tubulin normalized to total α-tubulin (*n* = 3). Data are mean ± s.e.m. Statistical significance was determined by one-way ANOVA with Tukey’s multiple-comparison test. ns, not significant; *P < 0.05, **P < 0.01, ****P < 0.0001.

**Figure S11.**
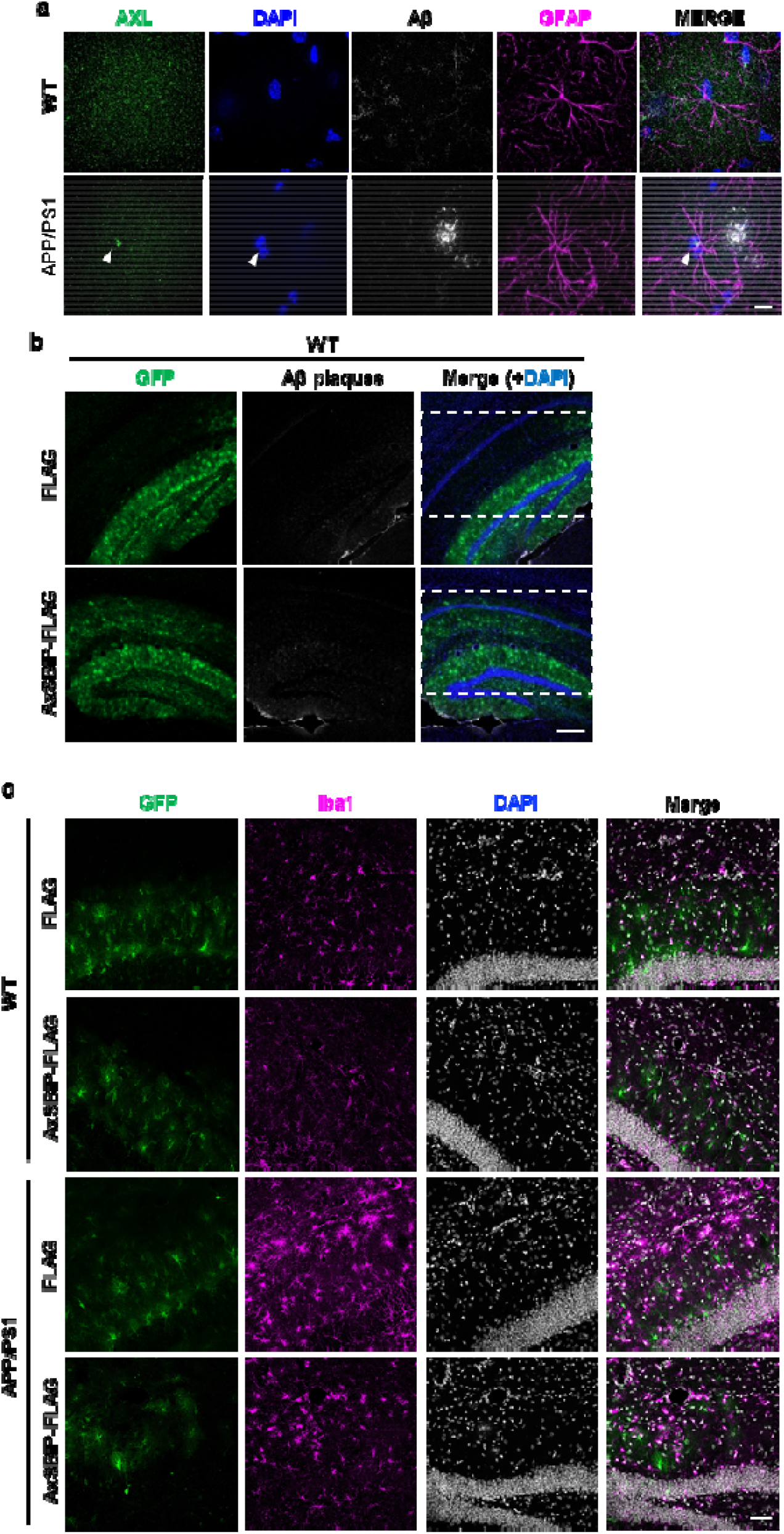
Nuclear localization of AXL in reactive astrocytes near Aβ plaques. **a** Immunofluorescence of brain sections from WT and APP/PS1 mice stained for AXL, GFAP and Aβ. Arrowheads indicate nuclear AXL signals in astrocytes adjacent to Aβ plaques. Scale bar, 10 µm. **b** Representative images of hippocampal sections from WT mice injected with the indicated viruses showing GFP-labelled cells and Aβ plaques. Scale bar, 200 µm. **c** Pseudocoloured representative confocal images of mouse hippocampus tissue stained to visualise AxSBiP/FLAG virus (green), Iba1 (magenta), and DAPI (white). Scale bar, 50 µm

## Notes

### Competing Interest Statement

The authors have declared no competing interest.

### Summary of Updates

No changes were made to the manuscript text, figures, or data in this revision."

